# Interference of mid-level sound statistics underlie human speech recognition sensitivity in natural noise

**DOI:** 10.1101/2024.02.13.579526

**Authors:** Alex C. Clonan, Xiu Zhai, Ian H. Stevenson, Monty A. Escabí

**Affiliations:** Electrical and Computer Engineering, University of Connecticut, Storrs, CT 06269; Biomedical Engineering, University of Connecticut, Storrs, CT 06269; Psychological Sciences, University of Connecticut, Storrs, CT 06269; Institute of Brain and Cognitive Sciences, University of Connecticut, Storrs, CT 06269; Biomedical Engineering, Wentworth Institute of Technology, Boston, MA 02115

## Abstract

Recognizing speech in noise, such as in a busy restaurant, is an essential cognitive skill where the task difficulty varies across environments and noise levels. Although there is growing evidence that the auditory system relies on statistical representations for perceiving ^1-5^ and coding^4,6-9^ natural sounds, it’s less clear how statistical cues and neural representations contribute to segregating speech in natural auditory scenes. We demonstrate that human listeners rely on mid-level statistics to segregate and recognize speech in environmental noise. Using natural backgrounds and variants with perturbed spectro-temporal statistics, we show that speech recognition accuracy at a fixed noise level varies extensively across natural backgrounds (0% to 100%). Furthermore, for each background the unique interference created by summary statistics can mask or unmask speech, thus hindering or improving speech recognition. To identify the neural coding strategy and statistical cues that influence accuracy, we developed *generalized perceptual regression*, a framework that links summary statistics from a neural model to word recognition accuracy. Whereas a peripheral cochlear model accounts for only 60% of perceptual variance, summary statistics from a mid-level auditory midbrain model accurately predicts single trial sensory judgments, accounting for more than 90% of the perceptual variance. Furthermore, perceptual weights from the regression framework identify which statistics and tuned neural filters are influential and how they impact recognition. Thus, perception of speech in natural backgrounds relies on a mid-level auditory representation involving interference of multiple summary statistics that impact recognition beneficially or detrimentally across natural background sounds.

**Significance Statement:** Recognizing speech in natural auditory scenes with competing talkers and environmental noise is a critical cognitive skill. Although normal listeners effortlessly perform this task, for instance in a crowded restaurant, it challenges individuals with hearing loss and our most sophisticated machine systems. We tested human participants listening to speech in natural noises with varied statistical characteristics and demonstrate that they rely on a statistical representation of sounds to segregate speech from environmental noise. Using a model of the auditory system, we then demonstrate that a brain inspired statistical representation of natural sounds accurately predicts human perceptual trends across wide range of natural backgrounds and noise levels and reveals key statistical features and neural computations underlying human abilities for this task.

## Introduction

Listening in competing noise is a central problem for both humans and animals given that isolated sounds rarely occur during real-world listening. The *cocktail party problem*, where conversational speech is relatively easily understood in a noisy social setting ^10^, is a classic example of the resilience of hearing in noisy scenarios. Humans often identify speech even when the power of environmental background noise exceeds the power of the speech signal (with signal-to-noise ratio, SNR, as low as ∼ -10 dB). However, noisy acoustic environments create substantial challenges for individuals with hearing loss ^11,12^ and even the most accurate automated speech recognition systems often fail in moderate levels of environmental noise. During such listening tasks, the original foreground and background sound pressure waveforms mix linearly in the air medium and, subsequently, nonlinear transformations throughout the auditory pathway create a complex neural interference pattern that must be segregated and interpreted by the brain.

The auditory neural pathways involved in segregating speech from environmental noise involves a hierarchy of processing that culminates in auditory cortex. Sounds are first decomposed by the cochlea into frequency components. Subsequently, sounds are decompose into detailed spectro-temporal modulations components in mid-level auditory structure (auditory brainstem and midbrain) ^13-18^ and slower and coarser modulation components in auditory thalamus and cortex ^15,19^. Human and animal studies further suggest that hierarchical computations produce a noise invariant representation in secondary cortices ^20-22^. Despite the important role of auditory cortex, there is compelling evidence that intermediate mid-level acoustic features, such as those represented subcortically in auditory midbrain^23^, are a primary source of interference that is necessary for separating speech from noise.

Studies also suggest that the brain may rely on mid-level statistical representations of natural sensory stimuli for coding and perception^3-5,23,24^. For instance, many natural sounds that often compete with speech, such as wind, fire, and crowd noise, can be described by summary statistics using a mid-level auditory model and the perceived realism of such sounds is highly contingent on unique summary statistics ^3-5^. Furthermore, direct measurements of neural response statistic from auditory midbrain (inferior colliculus) can be decoded to identify sound^4,6,7^ and these neural response statistics accurately predict human perception trends to natural sound textures ^4^. This supports the hypothesis that the brain relies on statistical information for recognizing and discriminating natural sounds. However, while such statistical representations may contribute towards defining the identity of sounds, it is less clear whether such statistical representations are directly involved in segregating and ultimately enabling speech recognition in competing environmental noise. It is known that spectro-temporal sound cues can impact sound segregation, for instance through energetic masking when the spectrum of a background sound overlaps and interferes with the spectrum of a foreground target^11,25^. And temporal fluctuations in power (amplitude modulations) can also create interference that can either obscure ^26-29^ or, in certain instances, facilitate the perception of a target sound ^30-32^. Although it is clear that such factors can impact speech perception with controlled laboratory background stimuli, it is less clear how arbitrary real-world background sounds influence and interfere with speech perception in natural auditory settings.

Recent advances in machine learning have enabled the develop of sensory processing models that are capable of accurately predicting neural activity^33,34^ and task optimized neural network models of hearing can produce human like behavior and adhere to known neural computations^35-39^, suggesting that auditory encoding mechanisms may be optimized for real world hearing tasks. Yet, models that are directly optimized and fitted to individual listeners behavior that can identify plausible neural coding or perceptual strategies during real-world listening are currently not available. Such models tailored to individual human listeners could help identify task specific neural mechanism and may be useful for developing real-world hearing diagnosis and optimizing prosthetic technologies.

We hypothesize that mid-level statistical cues in natural environmental sounds interfere with speech perception and that an auditory system inspired representation of these statistics explains behavioral performance for arbitrary background sounds. Using a diverse repertoire of natural background sounds and variants with perturbed spectrotemporal statistics to test human participants listening to speech in noise, we find that spectrum and modulation statistics in real world environmental sounds produce extensive background dependent differences in acoustic interference. These background dependent differences in accuracy span nearly 100% change in accuracy at a fixed SNR and can either improve or reduce speech recognition accuracy. We then develop *generalized perceptual regression* (GPR), a framework to relate statistical features from a biologically inspired auditory model to perceptual accuracy. GPR is an extension of the logistic regression classifier^40-42^ that incorporates a generalized neural model of sensory coding as input features and regresses against experimentally measured sensory decisions. This allows for studying complex natural behaviors with arbitrary high-dimensional natural sensory stimuli and testing different model representations. We demonstrate that a mid-level auditory model framework accurately predicts speech recognition behavior in arbitrary environmental noises and reveals key neural computations and multi-dimensional interference patterns that drive speech-in-noise recognition performance for natural background sounds. This provides an interpretable biologically grounded framework that links neural computations to perception, and which reveals a mid-level computational strategy for segregating speech from natural noise.

### Spectral and modulation statistics can mask and unmask speech

Human participants (n=18) listened to a series of randomly chosen digit triplets (e.g., five-nine-three; zero-seven-two, etc.) in the presence of natural and perturbed environmental noises and were asked to identify the digit sequence heard (Fig. 1A; see **Methods**). Here, ten natural environmental noises (e.g., crackling fire, speech babble, machine noise etc.) and white noise (as a control) were selected to encompass a wide range of natural sound spectra and modulation statistics (Fig. 1B, shown for 8-speaker babble; other sounds in **Figure 1-1 to 1-11**). In the first experiment, we examined how the spectrum and modulation statistics of natural background sounds separately influenced digit recognition, either beneficially or detrimentally. In addition to presenting original backgrounds (OR, Fig. 1B), we also presented perturbed environmental background noises where the contribution of the spectrum or modulation statistics towards listening in noise was isolated. In a first perturbation, the background waveforms were phase randomized (PR) which preserves the spectrum of each individual sound but randomizes and whitens the modulation statistics (PR, Fig. 1B). For these background sounds, only the background sound spectrum influenced digit recognition. These variants preserve the original sound spectrum and, thus, the unique timbre of the original backgrounds and are each perceived as spectrum shaped noise with no clear identity (**Audio 1-1**). In a second perturbation, the sound spectra of the backgrounds were equalized to follow a 1/f spectral profile ^4^ (spectrum equalization, SE). Since these SE backgrounds have identical spectra, only their individual modulation statistics influenced digit recognition across backgrounds (SE, Fig. 1B). Qualitatively, when listening to each of the SE variants, the original background can be easily identified (e.g., water, fire, street noise etc.), although all the SE sounds have similar timbre. That is, in contrast the PR variants, spectrum equalization preserves the spectrotemporal modulations that allow for the identification of sound category ^4^, so that fire sounds like fire and water sounds like water (**Audio 1-1**).

**Figure 1.**
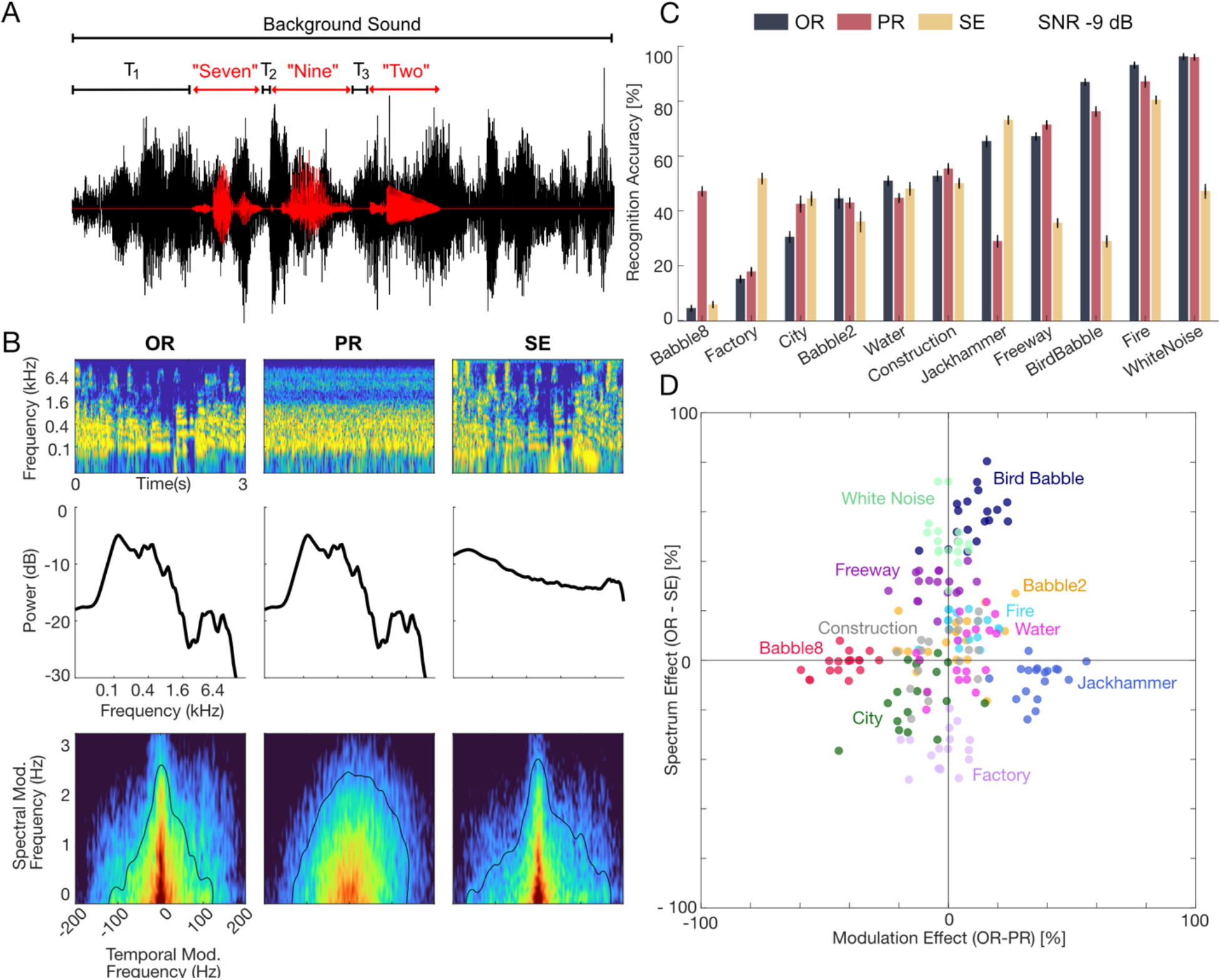
Experiment 1, The contribution of spectrum and modulation statistics to digit recognition in natural environmental noise. (A) Speech digits were delivered to human participants in competing natural background noises at -9 dB SNR (11 backgrounds in three configurations). The intervals between digits (T_1_, T_2_, T_3_) were randomly jittered to minimize learning and leveraging the word timing. Participants were required to identify the digit sequence (e.g., Seven-Nine-Two). (B) Each of the backgrounds was delivered in the original (OR), phase randomized (PR), and spectrum equalize (SE) configurations (illustrated for the speech babble; other sounds shown in **Figure 1-1 to 1-11**). Example audio excerpts are provided as part of the extended data (**Audio 1-1**). Cochleograms, cochlear power spectrum (cPS), and modulation power spectrum are shown in the 1^st^, 2^nd^, and 3^rd^ rows. The PR condition preserves the original spectrum of the sound (middle panel, 2^nd^ row) but whitens the modulation power (middle panel, 3^rd^ row). Alternately, the SE condition whitens the sound cPS, but retains much of the original modulation content. (C) Average results across n=18 participants show digit recognition accuracy in the different background sound conditions (rank ordered according to OR accuracy; OR=black, PR=red, SE=yellow). Removing the modulation (PR) or spectral (SE) cues can drastically change recognition accuracy, indicating that both cues contribute to masking. (D) Change in accuracy between OR and PR condition are indicative of a modulation masking effect while change between OR and SE are indicative of a spectral masking effect. Recognition accuracy shows clustering across participants for different backgrounds (color coded) indicating a predominant and sound dependent modulation (ordinate, e.g., babble-8, jackhammer) or spectral (abscissa, e.g., bird babble, white noise, or factory) masking effect. The spectral and/or modulation statistics in OR could either improve (e.g., jackhammer, bird babble) or reduce accuracy (e.g., factory, babble-8).

Digit recognition accuracy varied substantially for different background sounds and was strongly influenced by the unique spectrum and/or modulation statistics of each background (at a fixed Signal-to-Noise ratio, SNR, -9 dB). For the OR backgrounds, accuracy varied from near 0 (e.g., 4.4% for babble-8) to near 100% (e.g., 96.2% for white noise, fire, Fig. 1C, black) and perturbing the spectrum (SE, yellow) or modulation (PR, red) statistics could drastically modulate recognition accuracy. For instance, although the OR jackhammer noise showed moderately high accuracy (65.3%), the PR jackhammer noise (Fig. 1C, red) had substantially lower accuracy (28.9%). Since these two sounds have identical spectra, the modulation statistics in the OR jackhammer noise appear to have a beneficial unmasking effect on digit recognition. The spectrum-equalized jackhammer background (Fig. 1C; SE, yellow), which contains the original modulation statistics but has a whitened sound spectrum, had a high accuracy (73.1%). As for the OR condition, this suggests that original modulation statistics of the jackhammer sound had a beneficial impact on speech recognition accuracy. In stark contrast, the modulation statistics of eight speaker babble had a detrimental effect on digit recognition. Here, the OR eight speaker babble resulted in the lowest digit recognition accuracy of all backgrounds tested (4.4%). However, accuracy was substantially higher (47.1%) after whitening the modulation spectrum of the eight-speaker babble (PR condition). On the other hand, the SE variant which retained the modulation statistics resulted in a low accuracy (5.8%), similar to OR, implying that the modulation statistics strongly drove the perceptual interference. Across background sounds, both the PR and SE manipulations produced changes in accuracy that were highly dependent upon the specific background sound. These trends were conserved and highly consistent across participants (across participant *r* = 0.91 ± 0.04, mean ± sd) suggesting a common pattern of spectral and/or modulation interference.

Differences in the spectrum and modulation statistics of individual background sounds led to either masking or unmasking of speech digits (Fig. 1D). Here, changes in accuracy between OR and PR were used to assess modulation masking effects whereas changes in accuracy between OR and SE allowed us to isolate spectral masking effects. Backgrounds such as the freeway, bird babble, fire, and white noise had greater accuracy in the OR compared to the SE configuration, indicating that the original spectrum of these sounds had a beneficial effect. By comparison, for city and factory noise accuracy was higher for the SE compared to the OR condition, suggesting that the original sound spectra had a detrimental effect on digit recognition accuracy. Yet for other sounds, such as eight-speaker babble and the jackhammer noise, OR and SE background had similar accuracy, but OR and PR backgrounds differed, indicating effects due to modulation statistics. As with the spectral effect, the presence of the original modulation statistics could either be beneficial (e.g., jackhammer, PR<OR) or detrimental (e.g., babble-8, PR>OR) to accuracy. In each case, these effects were highly clustered across participants (Fig. 1D) suggesting that the unique spectrum or modulation statistics had a consistent impact on digit recognition.

In a second experiment, we further explored how background sound statistics contribute to recognition in noise by using texture synthesis ^3^ to systematically manipulate the background statistics. For these tests, we selected eight-speaker babble (n=10 participants) and jackhammer noise (n=6 participants) because they exhibit opposing accuracy trends driven by the modulation statistics (Fig. 1C, D). We generated synthetic variants of the babble and jackhammer backgrounds by sequentially incorporating summary statistics and repeated the digit recognition task with the synthetic backgrounds at SNRs -12 to -3 dB. Starting with a synthetic variant that strictly incorporated the spectrum statistics (Spec; similar to PR), backgrounds were sequentially manipulated to include marginal (+Mar), modulation power (+Mod), and correlation statistics (+Corr) (Fig. 2; A2-A4, B2-B4). Each of these summary statistics has unique structure that produces increasingly more realistic percepts as they are incorporated in the synthetic variants ^3,4,43-45^ (**Audio 2-1**). For the spectrum and marginal statistics (envelope mean and variance) both background sounds have similar structure (Fig. 2, A2 vs. B2)^43^. For the jackhammer, the modulation statistic (Fig. 2, A3; the sound power at specific frequency and modulation frequency combinations) is consistent with several audible components, particularly a high frequency (>∼2 kHz) dominant jackhammer sound with ∼20-30 Hz modulation frequency and a weaker low frequency engine noise with modulations > 10 Hz. By comparison, speech babble has greater energy for modulation frequencies < 10Hz (Fig. 2, B3) due to the slow rhythmic fluctuations present in speech ^44,45^ and also contains strong high frequency modulations (>∼100 Hz) that are generated by voice periodicity ^23^. Finally, the correlation statistics, which account for the strength of correlation between frequency channels, differ between the jackhammer and babble sound, with the jackhammer noise having somewhat broader pattern across frequency channels.

**Figure 2.**
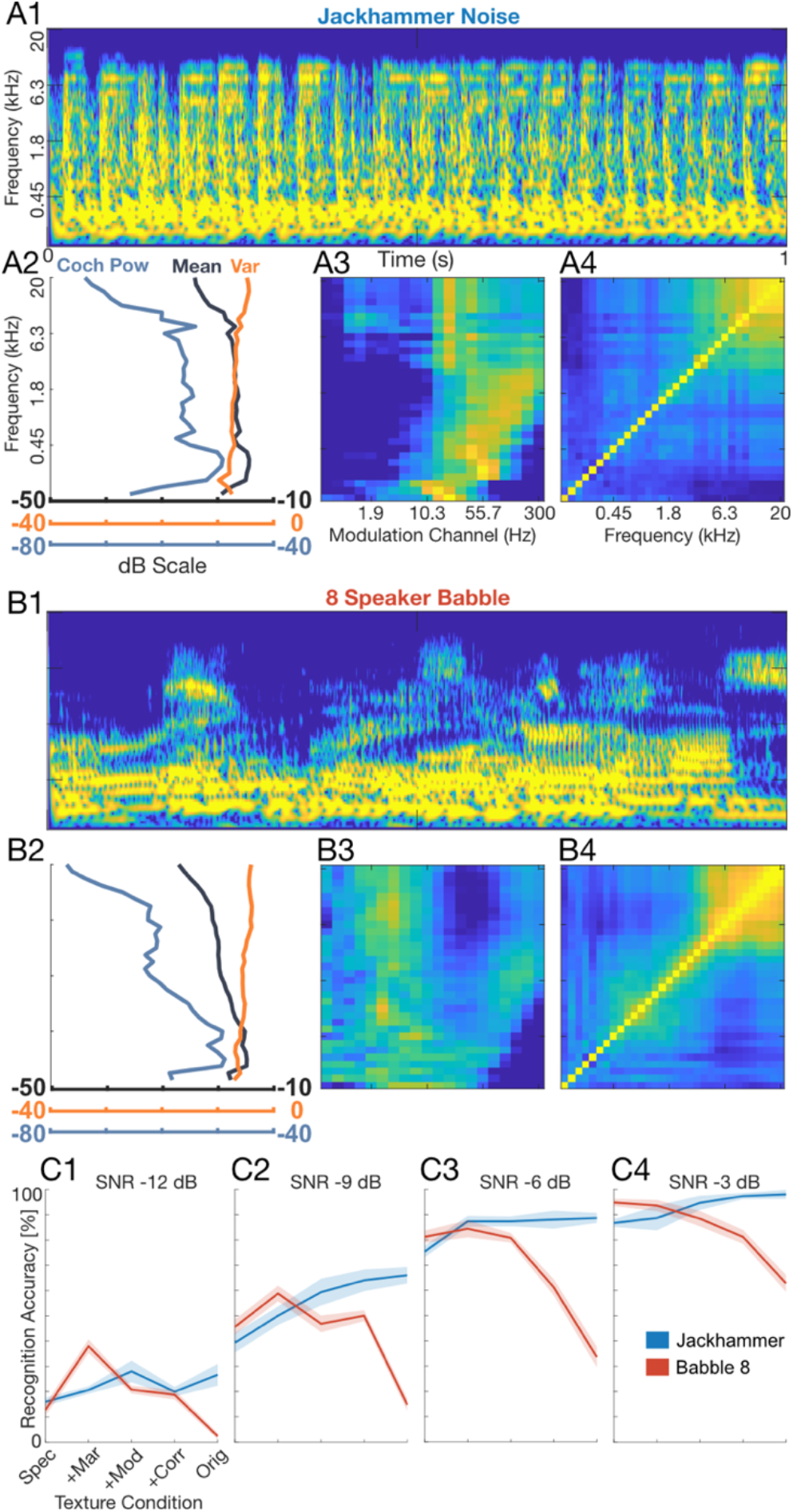
Experiment 2, Modulation summary statistics influence digit recognition accuracy. Sounds were delivered as in Experiment 1, but here, texture synthesis^3^ was used to manipulate individual summary statistics for the jackhammer (A1-A4) and babble-8 (B1-B4) backgrounds. Example audio excerpts are provided as part of the extended data (**Audio 2-1**). The cochleograms are shown in A1 and B1, along with the marginal (Mar; A2, B2), modulation power (Mod; A3, B3) and correlation (Corr; A4, B4) statistics which show unique structure for both sounds. Although Mar statistics are similar for both sounds (A2, B2), the Mod statistics show that speech contains extensive low frequency modulations (<10 Hz) along with fast temporal modulation arising from vocal fold vibration (B3; >55 Hz). Speech babble shows correlations across frequency channels (B4), but these are somewhat more compact than the jackhammer sound (A4). The jackhammer contains limited Mod power for low frequencies (<10 Hz) and contains a strong comodulated component at ∼20 Hz for frequencies > ∼2kHz and extensive correlations across frequencies (A4). (C1-C4) Digit recognition accuracy increases systematically with SNR, but also varies with the included summary statistics. For the speech babble (n=10 participants), accuracy decreases upon adding high-order summary statistics indicating that modulation structure in speech babble interferes with digit recognition. By comparison, for the jackhammer (n=6), including high-order summary statistics improves digit recognition accuracy indicating a beneficial unmasking. Error bars indicate SEM.

Digit recognition accuracy deteriorated upon adding high-order summary statistics to the synthetic eight-speaker babble backgrounds; however, and surprisingly, it improved upon adding summary statistics to the synthetic jackhammer (Fig. 2, C1-C4). For instance, at an SNR of -9 dB adding summary statistics to the synthetic eight-speaker babble reduced the digit recognition accuracy from 45.6% to 14.8%, while for jackhammer noise, adding statistics improved accuracy from 39.3% to 66.0% (Fig 2, C2). These effects appear to be largely independent of SNR, and the accuracy trends were well described by independent functions of SNR and summary statistics (separability index=0.98 ±0.01, mean±sd; see **Methods**). The improvement in recognition when adding statistics to the jackhammer noise may result from spectrally correlated fluctuations seen at ∼20 Hz (see **Figure 1-7**), since comodulation can in some instances unmask the perception of a target sound ^30,32^. Furthermore, the deterioration in recognition when adding statistics to the eight-speaker babble may reflect the fact that the spectrotemporal modulations in babble overlap closely with those of the speech digits. Once again, the perceptual trends were highly consistent across participants (across participant *r*=0.95 ±0.02, mean±sd) suggesting that a common set of perceptual strategies and cues influence recognition. Collectively, these results show that layering different levels of high-order acoustic structure in synthetic background variants can impact digit recognition, either beneficially or detrimentally.

### Low-dimensional interference of mid-level sound statistics predicts perceptual sensitivity

Using the experimental observations above, we next asked whether summary statistics derived from a biologically inspired auditory model can predict human recognition behavior. Recent studies support the view that hierarchical computations performed by peripheral and mid-level auditory structures are critical for the perception of environmental sounds ^4,5^. Here we propose a model based on the transformations that occur, first, at the cochlea and, subsequently, in mid-level auditory structures (inferior colliculus) ^23^ and use this model to predict speech in noise behavior with natural background sounds. Sounds are first decomposed by frequency, generating a cochleogram representation (Fig. 3A). Subsequently, sounds are decomposed through a set of modulation selective filters that model the decomposition of sounds into spectro-temporal components in auditory midbrain. While the time-average power spectrum distribution of the cochleogram (the cochlear power spectrum, cPS; Fig. 3, B2-D2) accounts for the distribution of sound power across frequencies, the midbrain modulation power spectrum (mMPS; Fig. 3, B3-D3) generates a multi-dimensional distribution of the sound power in the midbrain feature space. Here, sounds are represented as a function of frequency, and temporal and spectral modulation components. Speech digits and natural backgrounds not only differ in the spectral content, as seen in their cPS (Fig. 3, B2-D2), but they also have unique mMPS that can be similar (2-speaker babble, Fig. D3) or quite different (bird babble and jackhammer; Fig. 3, C3 and B3) from speech. Such differences in the power distributions between the foreground and background sounds has the potential to produce distinct patterns of acoustic interference. For instance, whereas the 2-speaker babble sound has high power that interferes with speech digits primarily at low frequencies (D2), the bird babble background sound interferes predominantly at high frequencies (B2), and the jackhammer sound interferes across the entire cochlear spectrum range (C2). While the cPS only accounts for frequency-based interference, the mMPS offers insight into how modulation components might interfere. For example, 2-speaker babble contains substantial power for temporal modulations within the vicinity of 4 Hz (D3, broadband 4 Hz peak visible at 0 cycles/octave spectral modulation) that have the potential to interfere with similar temporal modulations in the speech digit sequences. The jackhammer sound, by comparison, contains repetitive temporal fluctuations that produce a high-power band within the vicinity of ∼20-30 Hz temporal modulation and which are not present in the speech sequences (C3). These foregrounds and backgrounds also contain strong fine-structure fluctuations (visible for high temporal and low spectral modulations) that have potential for interfering at low frequencies (<1 kHz). The unique spectro-temporal modulations of each background have the potential to interfere with the modulations in speech to varying degrees and, thus, generate differences in recognition accuracy.

**Figure 3:**
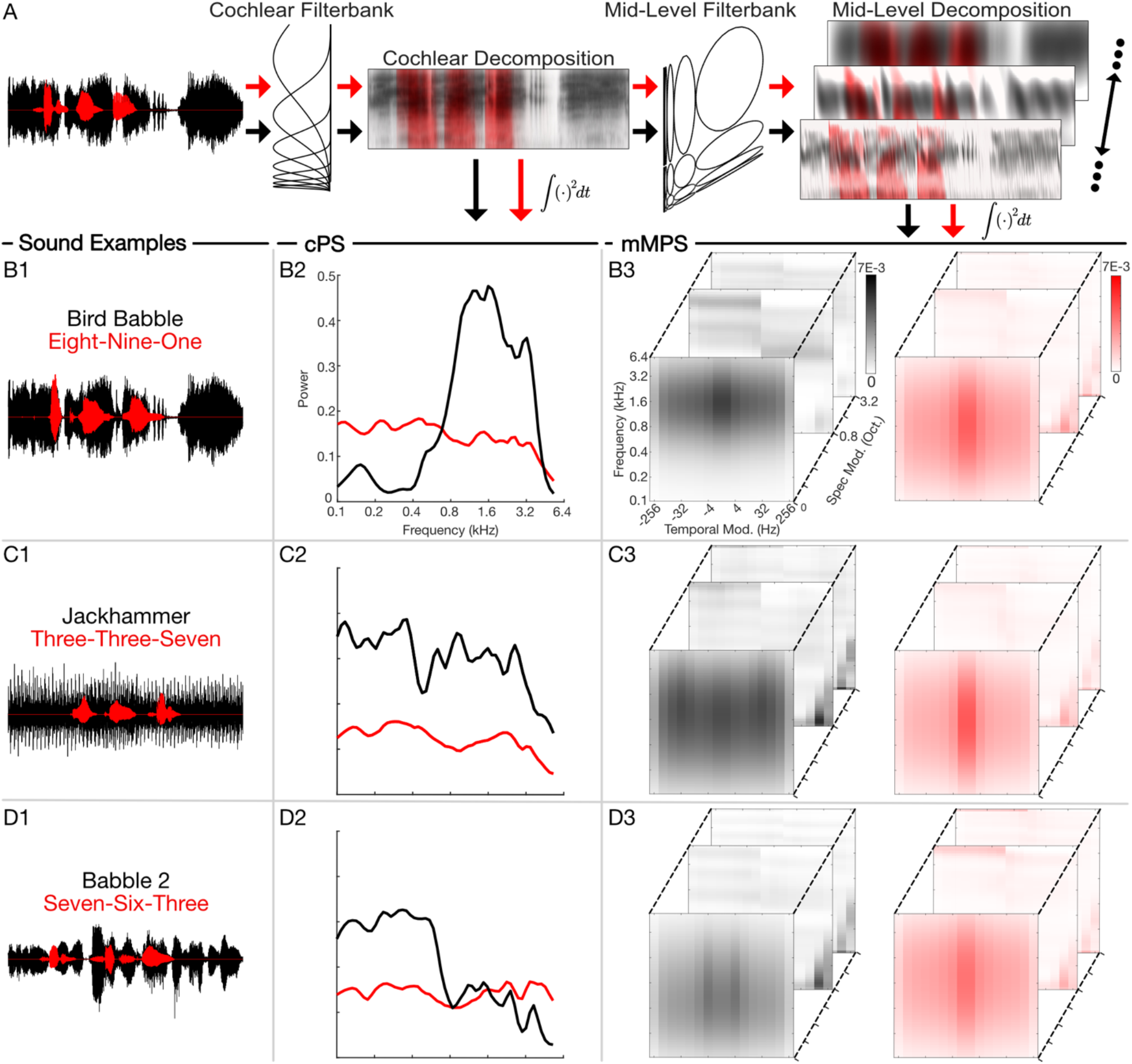
Cochlear and mid-level auditory feature representations. (A) Both, the foreground (red) and background (black) sounds are first decomposed into frequency components through a cochlear auditory model. These cochlear model outputs are then passed through a mid-level filterbank which further decomposes the sounds into time-varying spectral and temporal modulation components. (B-D) The time-averaged output power yields power spectrum distributions in either the cochlear or mid-level auditory space. The resulting cochlear power spectrum (cPS; B2-D2) and mid-level power spectrum (mMPS, B3-D3) are illustrated for three representative sound mixtures (B1-D1). Whereas the cPS only captures foreground and background sound power distributions as a function of frequency, the multi-dimensional mMPS distribution represents the sound as a function of frequency, spectral, and temporal modulation frequencies.

We model listener accuracy on single trials by fitting a GPR model. Within the GPR framework, the auditory model output summary statistics described above (Fig. 3) serve as input features to a logistic regression model (see **Methods** and Fig. 4). For the speech in noise paradigm, either the cPS or mMPS of both the foreground and background sounds serve as independent multi-dimensional auditory model inputs (Fig. 4 A1 and A2, respectively). Here, the foreground and background features (cPS or mMPS) adversarially compete to predict the perceptual finding, which could identify model-based summary statistics that drive masking or unmasking of speech. Comparing the result for cPS versus mMPS, on the other hand, can be used to identify how the cochlear or mid-level auditory representations differ and potentially explain (or don’t explain) the perceptual findings. We optimized the model parameters to predict the probability of “correct” (1) and “incorrect” (0) responses. The model coefficients identify natural background and foreground sound features that improve (positive coefficients) or reduce (negative coefficients) digit recognition accuracy. Based on our perceptual data, we expect that the cPS representation may be able to describe masking and unmasking related to spectral differences (e.g., OR vs SE), while the mMPS representation may be necessary to account for modulation effects (e.g., OR vs PR and differences based on high-order statistics).

**Figure 4:**
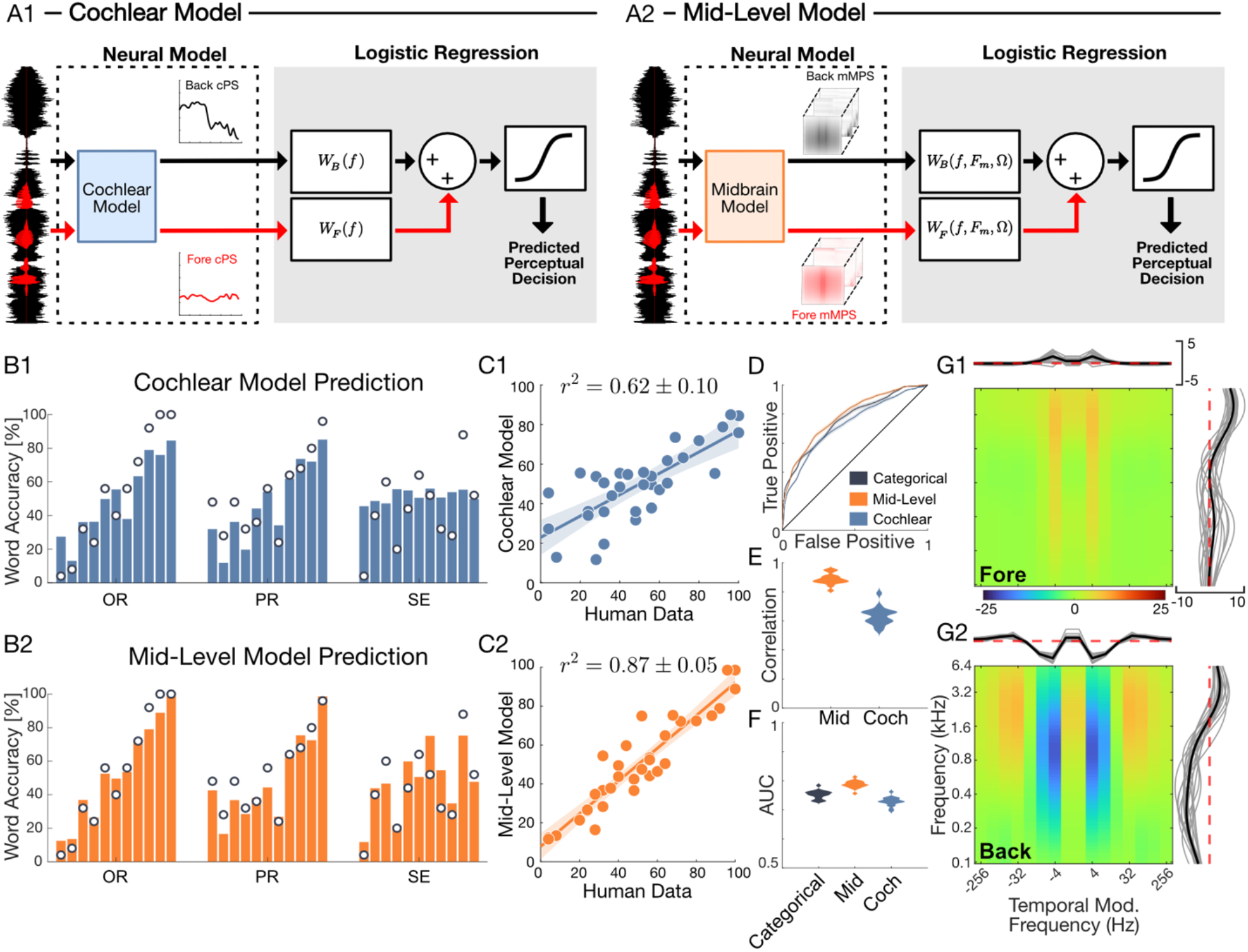
Mid-level auditory model predicts digit recognition accuracy in natural environmental noise. (A, B) The generalized perceptual regression (GPR) framework combines a cochlear (A1) or a mid-level (A2) model representation of natural sounds with logistic regression to predict single trial perceptual outcomes while listening to speech digits in noise. For both the cochlear and mid-level GPR model, the optimal regression weights were derived for each individual participant (**Figure 4-1**). Comparing digit recognition accuracy predictions for the cochlear (B1, C1) and mid-level (B2, C2) GPR models for a single participant show a vast improvement for the mid-level auditory representation. Bar plots (B1) show the cochlear model predicted accuracy for each of the three conditions of Experiment 1 (OR, PR and SE; background sounds are ordered as in Fig. 1C, from Babble-8 to White Noise). The predicted accuracies differ substantially from the actual measured accuracy for this participant (B1, superimposed dots; C1, *r*^2^ = 0.6 ± 0.1), especially for SE condition (B1, far right) where masking differences are attributed solely to the modulation content and the cochlear model produces a near constant prediction. (B2, C2) The mid-level model accounts for the modulation content of the sound and produces vastly improved predictions (C2, *r*^2^ = 0.87 ± 0.05). (D) The mid-level model ROC curve (participant from B1, B2) is higher than both the cochlear and categorical Bernoulli models. (E and F) Population results (n=18) confirm that the mid-level model predicts a higher proportion of the perceptual variance than the cochlear model (C; *r*^2^ = 0.89 ± 0.01 vs. *r*^2^ = 0.63 ± 0.01) and it outperforms both the cochlear model and categorical Bernoulli model on single-trial predictions as indicated by the higher AUC (F) and *d’* (Figure 4-2). (G1, G2) The mid-level model GPR regression weights can be interpreted as perceptual transfer functions. Here, the foreground (G1) and background (G2) model weights show antagonistic structure whereby specific frequencies and modulation components in the foreground and background can compete to improve or reduce digit recognition accuracy. The GPR weights of human listeners differed dramatically from those obtained for pretrained speech recognition neural networks (Figure 4-3), suggesting that humans employ a unique perceptual strategy. Notably, mid frequency (∼0.4-3.2 kHz) and slow modulation background components (∼4 Hz) tend to reduce accuracy (negative weights), while broadband speech driven components around ∼4 Hz show positive weights that can improve digit recognition accuracy for human listeners.

We first tested whether interference in the cochlear auditory space can account for differences in digit recognition. Here the cPS of both the foreground and background serve as independent inputs to a GPR model designed to predict recognition accuracy (Fig. 4, A1). Conceptually, the background and foreground sound power at each frequency are independently weighted by the model so that the foreground and background confer an opponent-weighted contribution to the predicted perceptual accuracy. For instance, when the background power at a critical frequency exceeds that of the foreground power, the predicted accuracy should be reduced. By comparison, increasing the foreground power at the same frequency should produce an increase in predicted accuracy. That is, the foreground and background cPS should produce an interference pattern with antagonistic influence on the predicted accuracy. To minimize the complexity of the model and possibility of overfitting, we used a low-dimensional representation of the cPS (20 dimensions; see **Methods**) and the model weights were optimized and predictions cross-validated (25-fold) for each participant (**Figure 4-1**). Results for a single participant demonstrate that the cochlear model has some predictive power and accounts for coarse sound dependent trends. The average accuracy predicted by the model for each background condition has a strong correlation with the observed accuracy of each condition (Fig. 4B1, C1; single participant *r*^2^ = 0.6 ± 0.1, mean ± sem). The model captures coarse trends for the OR and PR conditions (Fig. 4, B1; left and middle panels; dots are the participant accuracies) and, compared to the behavioral data, it has a relatively narrow output dynamic range (12-85 % versus 0-100 % for the behavior). Furthermore, although cochlear model accurately predicts the PR condition (*r*^2^ = 0.84 ± 0.14, mean ± sem) it fails to predict changes in accuracy when the sound spectra are equalized (SE condition; Fig. 4B1, right panel; *r*^2^ = 0.07 ± 0.32, mean ± sem) and it does not differentiate OR and PR responses, since it produces similar predictions for both (Fig. 4B1, left and middle; since OR and PR sounds have identical cPS).

We next tested whether multi-dimensional interference in a mid-level auditory space better explains the perceptual findings (Fig. 4 A2). Here, a low-dimensional representation of the mMPS is again used as input to the GPR model, which limits the model complexity and minimizes overfitting (optimal solution, 24 dimensions; see **Methods**). For the same participant, there is a marked improvement in the predicted accuracy for the mMPS input (Fig. 4, B2, C2; single participant *r*^2^ = 0.87 ± 0.06, mean ± sem) compared to the cPS. In addition to accurately predicting the OR and PR conditions, the mMPS input also captures modulation dependent changes in accuracy for the SE condition (Fig. 4B2, right panel). Across the sample of participants (n=18), the mid-level model outperforms the cochlear model and explains nearly 90% of the behavioral variance (Fig. 4D-F). The predictive power of the mid-level model is significantly higher than the cochlear model (Fig. 4E; average *r*^2^ = 0.89 ±0.01 vs. *r*^2^ = 0.63 ±0.01, mean ± sem; Fisher z − tra*n*sfor*m* paired t − test, t(17) = 20, p < 10^−12^) and this was true even when the mid-level model was constrained to the same dimensionality as the cochlear model (20 dimensions, average *r*^2^ = 0.85 ± 0.01, mean ± sem; *F*isher z − transform paired t − test, t(17) = 15.7, p < 10^−10^).

GPR also provides accuracy predictions for individual trials, and the mid-level representation can account for accuracy differences, not just across conditions (e.g., white noise OR vs babble-8 OR), but also within each condition (e.g., two different trials of babble-8 OR noise). Subtle differences between specific foregrounds and differences between multiple instances of the same background sound may have predictable effects on accuracy that are not apparent when all the trials within a condition are averaged. As a reference, we fit a *category-specific Bernoulli* model which uses the average accuracy of each sound condition on training data to predict test trials (33 parameters for OR, PR, and SE of each of the 11 background sounds). While this category-specific model lacks trial-specific information (null hypothesis), the mid-level model uses acoustic information from each trial, which could potentially account for trial specific changes in accuracy. For a single participant, the mid-level model receiver operating characteristic (ROC) curve is above those of both the category-specific and cochlear models (Fig. 4D), suggesting an improvement in single trial prediction quality across all trials. Across participants, the area under the ROC curve (AUC) for the mid-level model (Fig. 4F; 0.79 ± 0.01, mean ± sd) was significantly larger than the AUC of the category-specific and cochlear models (Fig. 4F; 0.75 ± 0.01 and 0.73 ± 0.02, mean ± sd; t(17)=12, p<10^−9^ and t(17)=16.5, p<10^−11^, paired t-test). Furthermore, while the category specific Bernoulli model has a sensitivity index (*d’*) of 0 for single-trial predictions, the mid-level model *d’* was significantly larger (participant average *d’*=0.31 ± 0.03, mean±sem; t-test, t(17)=11.6, p<10^−8^; **Figure 4-2**). Thus, the mid-level model not only outperforms the cochlear model in predicting differences between background conditions, but it also accounts for trial-by-trial changes in perceptual accuracy within background conditions.

To further explore the source of acoustic interference between speech and natural background sounds, we inspected the model weights and projected them onto frequency and temporal modulation dimensions of the mid-level auditory model. The foreground (Fig. 4G1) and background (Fig. 4G2) weights have an antagonistic relationship that depends both on the sound frequency and modulation content. For the background, model weights show strong broadband negativity centered at ∼1 kHz and very slow temporal modulation frequencies (∼4-8 Hz; Fig. 4G2). Backgrounds that contain substantial power within this modulation band, as seen for 2-speaker babble (Fig. 3 D3, ∼4-8 Hz), interfere more strongly with speech leading to reduced recognition accuracy. By comparison, at high frequencies (>0.8 kHz) and for temporal modulations between ∼16-64 Hz, the background model weights are positive, suggesting that background energy for these components enhances recognition accuracy. This range likely reflects a beneficial interference consistent with comodulation masking release ^30-32,46^, as observed for the jackhammer which contains strong comodulated fluctuations that overlap this acoustic range (Fig. 3 C3) and which has reduce accuracy for the PR manipulation (Fig. 2; accuracy enhancement in Fig. 3 when statistics are included). For the foreground sound, the model weights show a positive component within the vicinity of 4 Hz temporal modulation that opposes the negative component in the background and increases accuracy for foreground speech sequences with strong temporal modulations in this range. For reference, we compared human listeners results against those from two pretrained time delay speech recognition neural networks to identify strategies for listening in natural noise that are unique to human listeners (Figure 4-3). Compared to humans, both networks exhibited lower accuracy when tested across different SNR (Figure 4-3, A1 and A2) or at a single SNR (−9dB, Figure 4-3, B1 and B2). Furthermore, while the transfer function of both networks exhibited spectro-temporal components that masked or unmasked speech (Figure 4-3, C1, C2 and D; blue and red hot-spots), the frequency and modulation components that drove interference in these two networks differed extensively from those of human listeners. This suggests that humans rely a unique set of spectrotemporal statistics for segregating speech from noise and that background statistics can either facilitate or hinder word recognition.

### SNR contributes to speech recognition independently of spectro-temporal features

Beyond the statistical structure of natural sounds, background sound power can vary extensively in real world listening environments, which influences the listening SNR and ultimately the perceptual difficulty. Although lower SNRs clearly make it more difficult to hear and identify sounds, how the SNR interacts with speech perception for specific natural background sounds and their related statistics is unclear. Here we explored how SNR impacts speech recognition across natural background sound conditions.

Nine participants listened to digit triplets embedded in the non-perturbed, natural backgrounds (OR) at variable SNRs (−18 to 0 dB) (see **Methods**). The paradigm is challenging (average accuracy 55%) but covers a broad range of perceptual accuracies across both SNR and backgrounds, extending from very easy (e.g., 100 % for white noise at SNR=0 dB) to extremely difficult (e.g., 0 % for babble-8 at SNR=-18). As shown for a single participant, digit recognition accuracy increased with increasing SNR for all background sounds tested in a sigmoidal fashion with SNR (Fig. 5A). However, the intercept (inflection point of the sigmoid) varied across backgrounds, consistent with the fact that the spectrum and modulation statistics of each background drastically impact listening difficulty even at a constant SNR (as in Fig. 1, -9 dB). To quantify these differences, we estimated the SNR where digit recognition accuracy is 50% for each background (the logistic midpoint). Across participants (n=9, Fig. 5E), backgrounds that interfered strongly with foreground speech, such as the eight-speaker babble, had a high logistic midpoint SNR (∼ –3 dB for 50% accuracy). By comparison, backgrounds that do not extensively overlap with the foreground speech in either the spectrum and/or modulation statistics, such as bird babble, interfered less with speech perception and thus showed a much lower logistic midpoint SNR (∼ – 16 dB at 50% accuracy). Thus, to maintain 50% accuracy across background sounds, the SNR needs to be adjusted by as much as a 13 dBs. These general trends were highly consistent across listeners as noted by the relatively narrow standard deviation of midpoint distributions (Fig. 5E) and high inter-participant correlation (*r* = 0.95 ± 0.005, mean±sem).

**Figure 5.**
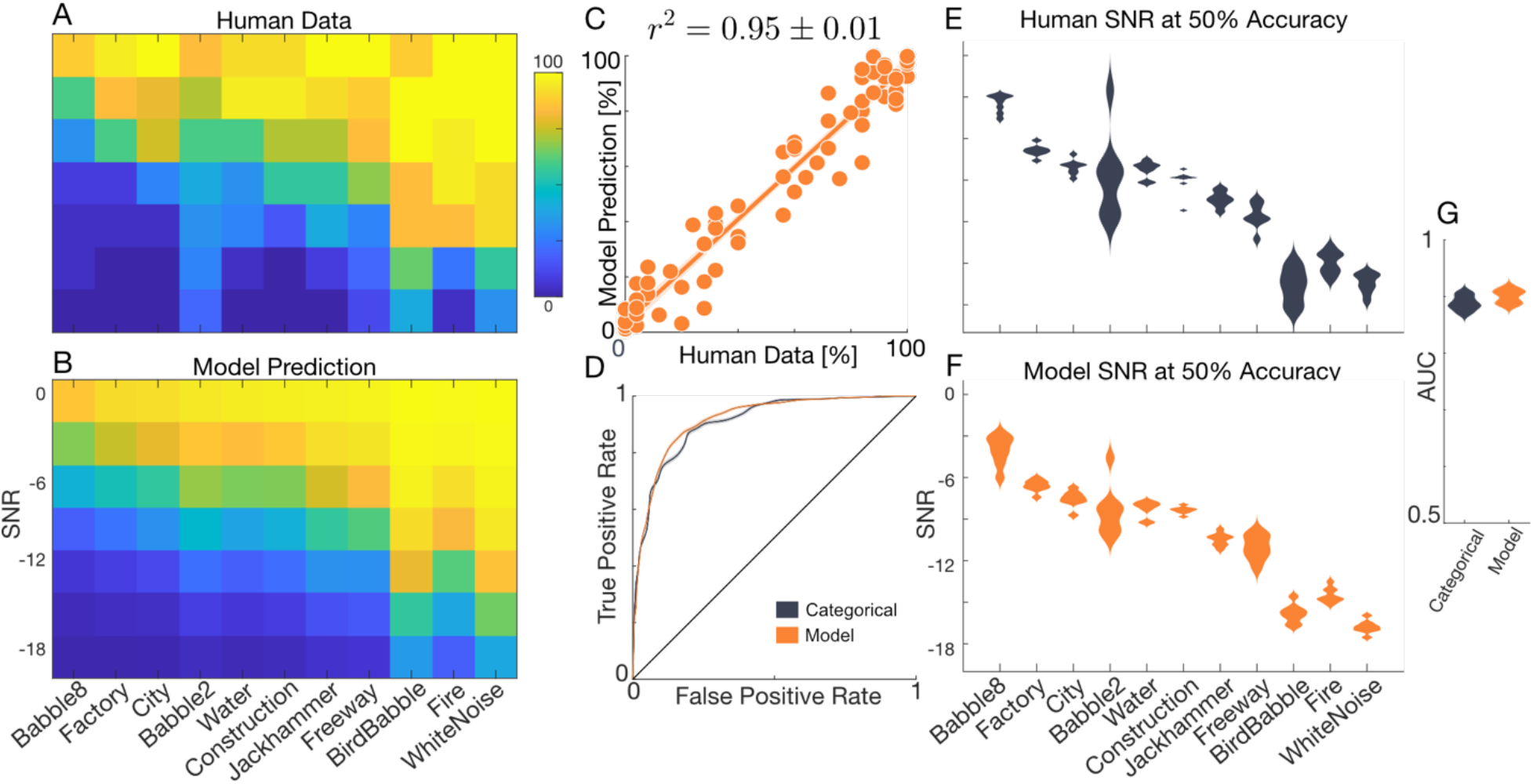
Experiment 3, SNR influences recognition independently of summary statistics. (A) Average accuracy for one participant is shown as a function of SNR and each of the eleven OR backgrounds. (B) Predictions using a modified mid-level model with SNR as an independent input variable accurately predict the participants digit recognition accuracy. (C) Scatter plot comparing the actual (A) versus the modified mid-level model predicted digit recognition accuracy (B). Across participants, the modified mid-level model explains more than 90% of the behavioral variance (*r*^2^ = 0.93 ± 0.005, mean ± sem). (D) ROC curve for the modified mid-level and a 77-parameter categorical Bernoulli model for the same participant as panels A and B. The distribution of logistic midpoints across nine participants and background sound conditions (E) is accurately predicted by the modified mid-level model (F). The AUC for the modified mid-level and categorical Bernoulli models (G; AUC=0.90 ± 0.005 vs. 0.89 ± 0.005; mean ± sd).

To determine whether the summary statistics of the background sound and SNR contribute independently to perceptual accuracy, we computed a separability index for the SNR across background trends (see **Methods**). The separability index was near 1 (0.93 for participant of Fig. 5A; population average=0.93 ± 0.01, mean ± sd) consistent with the hypothesis that SNR makes an independent contribution to masking. We additionally modified the GPR model by incorporating SNR as an independent input variable. The model incorporates the same spectrotemporal features used for the mid-level representation and adds a single independent regression weight to account for the SNR dependency of all the backgrounds (see **Methods**). This model accurately reconstructs digit recognition accuracy trends across backgrounds and SNRs for a single participant (Fig. 5B, C; *r*^2^ = 0.95 ± 0.01, mean ± sem). ROC analysis further confirms that, for this participant, the mid-level model is comparable to a category-specific Bernoulli model (Fig. 5D; AUC=0.92 vs. 0.91) in predicting single trial outcomes across all conditions. Across participants, the model predicts more than 90% of the perceptual variance (*r*^2^ = 0.93 ± 0.005, mean ± sem), accurately replicates the logistic-midpoint distributions (Fig. 5 E vs. F), and slightly outperforms the categorical model (Fig. 5G; AUC=0.90 ± 0.005 vs. 0.89 ± 0.005; mean ± sd; paired t-test, t(8)=8.3, p<5×10^−5^). Together, these results suggest that spectrotemporal sound features and SNR make independent contributions to digit recognition accuracy and demonstrate that a modified mid-level neural model can accurately explain human performance at across noise levels.

## Discussion

Our results suggest that speech recognition in competing natural background sounds involves the interference of mid-level summary statistics. For natural sounds, the spectrum and modulation statistics can be modeled through a biologically inspired auditory model, which accounts for both the spectral decomposition of the cochlea and the modulation decomposition in mid-level auditory centers ^23^. A low-dimensional representation of these mid-level sound statistics accounts for and explains the diverse types and amount of perceptual interference created by real-world environmental sounds. Importantly, our behavioral and modeling results suggest that spectrum and modulation statistics influence recognition independently of the overall noise level (SNR) and that complex masking and unmasking phenomena can be described through interpretable perceptual transfer functions.

### Using GPR to identify neural computations underlying perception

There is a long history of cognitive models that attempt to explain human psychometric performance yet, traditionally, these have been applied to the perception of relatively simple synthetic stimuli and their parameters (e.g., visual or auditory). Here we develop a modeling framework, GPR, which addresses several key obstacles that have hindered progress in linking neural models with models of sensory choice. First, GPR is an embedded model framework that combines a generalized neural model with logistic regression ^40-42^. The neural model compartment is interchangeable, which allows for testing and identifying different neural mechanisms that explain perceptual judgments. For speech-in-noise task, incorporating second-order mid-level transformations known to exist in the auditory midbrain helps explain modulation-based interference of natural sounds, which as we demonstrate is not possible with a cochlear model. Second, while logistic regression has been previously used to explain sensory choices, its usage is generally limited to varying a few relatively simple physical stimulus dimensions (e.g., intensity, contrast, frequency etc.) or model-based features^47,48^ that are used as regressor variables. The GPR model framework extends these methods by utilizing the transformed sensory inputs through a neural model as the regressor variables. When combined with dimensionality reduction, this enables the use of complex multi-dimensional natural stimuli in generalized perceptual tasks, such as the speech-in-noise task performed here. As shown, natural sounds mixtures span multiple entangled dimensions that interfere and jointly influence speech recognition performance. Finally, GPR can identify multiple model-based neural dimensions and statistics that underly perceptual trends and provides interpretable model transfer functions that can inform cognitive and neural processing theories. When combined with mid-level auditory representation of natural sounds^23^, the GPR approach identifies salient model features that impact, beneficially or detrimentally, speech recognition in real world listening scenarios. This is similar to conventional regression methods that has been used to identify auditory features that correlate with perceptual sound quality^49^. GPR is in part motivated by conventional regression methods used to estimate neural receptive fields, which relate sensory features to neural spikes (0s or 1s indicating presence or absence of a neural response) for natural sensory stimuli ^50,51^. Here, in lieu of neural spike trains, GPR uses single trial perceptual judgments (0s or 1s, indicating correct and incorrect decisions). Thus, future studies may be able to identify acoustic features driving both neural representations and perception in animal models by combining receptive field mapping methods and the GPR framework, thus linking neural coding of natural sensory stimuli to the overall perceptual strategy.

Recently, task-optimized deep neural network models have been developed that can recognize or categorize natural sensory stimuli and, interestingly, these networks can exhibit human like performance in both auditory and visual tasks ^36,52^. For listening tasks, such models can recognize individual words in competing noise with human-like accuracy despite the fact that they are not trained directly on human perceptual data^36^. Yet, these models often underperform in real environments or when exposed to novel datasets. Interpreting these complex neural network models and their relationships to cognitive and brain processing mechanisms remains challenging since neural networks are extremely nonlinear systems that typically contain tens-of-thousands to millions of parameters. This contrasts the GPR framework, whereby the model weights are low-dimensional and highly interpretable and can be viewed as perceptual transfer functions that map the model-based acoustic features to behavioral accuracy. Indeed, and as demonstrated for matched perceptual tasks, neural networks can be fit and projected onto the GPR which allows for comparing the computational strategies of these neural networks with human performance (**Figure 4-3**). As seen, human performance not only exceeds that of two pretrained neural networks, but their GPR weights are quite different from those of human listeners indicating that they use different features to segregate speech from noise. This is consistent with recent observations showing that neural networks models exhibit task-specific invariances that diverge from those of human listeners^53^ suggesting that their computational strategies can be quite different. This demonstrates how human and machine perception differ and how human perception is uniquely adapted to resolve speech from noise.

### Summary statistics and neural computations underlying speech in noise performance

The mid-level auditory model used in our GPR framework is a simplified representation of the computations carried out by the auditory system and the mid-level neural statistics contributing to recognition in noise. The principal auditory midbrain nucleus, the inferior colliculus, is a likely candidate for representing such acoustic information since neural response statistics from auditory midbrain covary with natural sound statistics and accurately predict human perception of sound textures ^4^. This contrast responses from auditory cortex to sound textures which show weak tuning to changes in summary statistics ^7,8^. Thus, our primary conclusion and claim is that average statistics from features homologous to those encoded in mid-level auditory system can explain the multi-dimensional interference between speech and natural background sounds. As shown, spectro-temporal modulation statistics of the competing background sound strongly impact recognition accuracy and the comparison between the cochlear and mid-level model predictions demonstrate a clear improvement for the mid-level representation. Whereas the cochlear model relies strictly on frequency-based interference to predict behavioral outcomes, the modulation-based interference in the mid-level model more accurately predicts perceptual outcomes.

The perceptual transfer functions derived using GPR directly link peripheral and mid-level statistical feature from the auditory model to perceptual accuracy. This provides an interpretable framework that identifies the specific sound cues and the perceptual strategy underlying speech in noise recognition performance. The mid-level model transfer functions suggest that slow fluctuations in background sounds (∼4 Hz) are a critical source of interference that competes with comparable fluctuations in speech. This is consistent with prior observations indicating that cortical activity for slowly varying non-stationary sounds is stronger than for stationary backgrounds ^21^ and may thus interfere more strongly with speech. The mid-level transfer functions also identify frequency and modulation specific enhancements in recognition accuracy (frequencies >0.8 kHz and temporal modulations ∼16-64 Hz) that is consistent with comodulation masking release phenomena ^30-32,46^.

The experimental paradigm employed was intentionally designed to perturb and identify the spectrotemporal sound statistics driving interference of speech and does not allow for linguistic factors that also play a role during conversational speech perception. For instance, lexical and semantic top-down signaling influence speech perception during continuous speech ^54-56^. Our perceptual paradigm intentionally avoids such influence by using randomly ordered digits, at variable onsets, from randomly selected talkers, thus focusing exclusively on the acoustic features underlying masking. Our model also does not account for attention and informational factors such as varying degrees of stimulus uncertainty ^57,58^ which also influence speech recognition in noise. By adding temporal structure and additional linguistic variables to the GPR model framework, future studies may be able to account for continuous speech recognition in natural environmental noise under a wider range of scenarios.

### Summary

The findings demonstrate that recognition accuracy in a complex real world behavioral task can be mapped onto a low-dimensional and quasi-linear representation of a biologically inspired model feature space. This generalized model framework yields an interpretable and plausible neural inspired representation that provides insights into the features and neural computations that underlie perceptual judgments for natural sensory signals. GPR can theoretically be applied to a variety of behavioral tasks across various sensory modalities and has the potential to be extended to animal behavior and model performance evaluation. For the speech in noise task, GPR reveals multi-dimensional natural sound features and transformations underlying masking and unmasking of speech by natural background sounds. Since this framework can make predictions for arbitrary background and foregrounds sounds, it may be useful for studying a wide range of complex real world listening scenarios and may allow for rapid clinical diagnostics that can identify real world listening deficits.

## Materials and Methods

### Human Psychoacoustics

All procedures were approved by the University of Connecticut Institutional Review Board. Participants were provided verbal and written description of the experiment procedures and study rationale prior to consenting to participate in the experiments. We recruited male (n=9) and female (n=18) native English speaker participants ages 18-43 with normal hearing sensitivity (>20 dB HL for left and right ears; 0.5, 2, 8 kHz, tested with MA-42 audiometer (Maico Diagnostics Co.). Experiments were carried out in a sound shield room (Industrial Acoustics Co.) at the University of Connecticut, Storrs campus. Sounds were delivered at 44.1 kHz sampling rate using an Fireface UC digital audio interface (RME Audio Inc.), amplified through a BHA-1 headphone amplifier (Bryston Ltd.), and delivered through HD 820 headphones (Sennheiser Co.) at 70 dB SPL.

Experimental sounds consisted of speech digits from zero to nine vocalized by eight male talkers (TI46 LDC Corpus ^59^). Digits were delivered in triplet sequences with random inter-digit intervals (Fig. 1A, *T*_1_ - *T*_3_), randomly chosen digits and talkers in the presence of a competing natural background noises (Fig. 1A). The silent intervals (*T*_1_ - *T*_3_) were not included in the estimation of the speech power when computing SNR. All sounds lasted 3.5 seconds duration and were gated with a 100 ms b-spline ramp to avoid broadband transients. Eleven environmental noises were selected to represent sounds widely encountered in nature (two- and eight-speaker babble, bird-babble, factory, city, construction, jackhammer, freeway, water, fire as well as white noise as a control) and which encompass a diverse range of spectrum and modulation statistics. For each of the experiments below, all the original and perturbed sound variants were delivered over 25 trials in block randomized order. Following each sound trial, participants were required to type the digit sequence heard on a keyboard.

#### Experiment 1

This experiment was carried out in n=18 participants. The experimental sounds are designed to assess how the spectrum and modulation statistics of natural environmental backgrounds interfere with speech recognition. In addition to delivering digit triplets in the original backgrounds (OR), which contains the original spectrum and modulation statistics, we delivered digits in phase randomized (PR) background variants which preserve the sound magnitude spectrum while randomizing the phase spectrum, thus distorting (whitening) the modulation statistics. These sounds are equivalent to spectrum shaped white noise. We also delivered digits in spectrum equalized (SE) background variants ^4^. Here the background spectrum is whitened but the modulation statistics are preserved. All sounds were delivered at an SNR of -9 dB.

#### Experiment 2

This experiment explores how individual summary statistics of natural background sounds interfere with and influence speech recognition. Here, we used jackhammer (n=6 participants) and eight-speaker babble (n=10 participants) backgrounds since these sounds exhibited opposing recognition trends driven by the modulation and spectrum statistics of the background (Fig. 1C, D). Texture synthesis ^3^ was used to generate synthetic background variants that sequentially incorporate the 1) spectrum (Spec), 2) marginal (+Mar), 3) modulation power (+Mod), and 4) correlation (+Corr) summary statistics of each background. Sequentially adding summary statistics to the synthetic variants creates backgrounds that are perceptually more realistic and more closely match the statistics of the original background ^3^. SNRs were varied between -12 to -3 dB (3 dB steps).

#### Experiment 3

In this final experiment, we tested whether summary statistics from the OR backgrounds influence speech recognition independently of the noise level (SNR). The original backgrounds used for Experiment 1, where delivered in the OR configuration at multiple SNRs. For one group of participants (n=5 participants) sounds were delivered at -15, 12, -9, -6, -3 dB SNR. For n=4 participants, the range was extended to also include -18 and 0 dB.

In experiments 2 and 3, where the listening SNR was varied, we evaluated whether SNR influenced perceptual accuracy independently of the sound statistics. Here, singular value decomposition was first used to determine whether the response accuracy matrices are separable functions of the included background sound statistics and SNR (for experiment 2; background sound vs. SNR, for experiment 3). The separability index was defined as the fractional power explained by the first singular value, 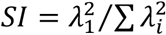. SI values near 1 indicate that the response accuracy matrix is separable, so that SNR influences accuracy independently of the background statistics. Secondly, in addition to considering spectro-temporal features of the background sound as variables that can predict perceptual accuracy (for the model-based approach below), we also included SNR as an independent explanatory variable into the GPR model framework. This approach provides a secondary model-based assessment of independence between SNR and spectro-temporal acoustic features.

### Peripheral and Mid-Level Acoustic Features

We used a two-stage hierarchical auditory model representation of the auditory system to derive time-average summary statistics for both the foreground and background sounds and these time-averaged statistical features are subsequently used to predict human speech recognition in noise behavior. The auditory model has been described previously ^23^ and is freely available for non-commercial use ^60^. Thus, only necessary details are outlined below.^23^ and is freely available for non-commercial use ^60^. Thus, only necessary details are outlined below.

Both the foreground and background sounds for each trial in Experiments 1 and 2 were first decomposed through peripheral filterbank that models the frequency decomposition, envelope extraction, and nonlinear rectification of the cochlea. At this stage, sounds are represented by a cochlear spectrogram, *S*(*t, X*_*l*_), where *X*_*l*_ represents the octave frequency of the *l*^th^ channel (57 filters between 0.1-5.3 kHz) and *t* is the time variable. The time-averaged cochlear power spectrum is then estimated as:

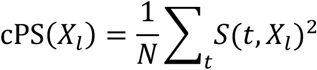

where *N* is the number of time-samples. The cPS summary statistics accounts for the spectral power distribution as represented through a cochlear model. It is subsequently used to assess the contribution of spectrum-based interference between the foreground speech and background.

Next, to represent the sounds in a mid-level auditory feature space, the foreground and background cochlear spectrograms are passed through a set of modulation selective filters that model the decomposition of sound into spectrotemporal modulation component in auditory midbrain. Here, each sound is represented as a multi-dimensional midbrainogram, *S*(*t, X*_*l*_, *f*_*m*_, Ω_*n*_), where *f*_*m*_ is the temporal modulation frequency for the *m*^th^ channel (filter modulation freq. of 0, ±2 to ±256 Hz in octave steps) and Ω_*n*_ is the spectral modulation frequency of the *n*^th^ channel (0 and 0.1 to 3.2 cycles/oct in octave steps). The midbrain model derived modulation power spectrum is then computed as

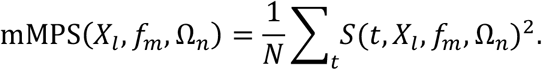

The mMPS summary statistics represents the sound power distribution jointly across frequency channels and combinations of spectral and temporal modulation channels, as represented through an auditory midbrain decomposition. This multidimensional sound representation is used below to model the interference between background and foreground speech sounds across spectral and modulation components in order to predict speech recognition in noise behavior.

### Modeling Acoustic Interference Through Generalized Perceptual Regression

We developed *generalized perceptual regression*, a framework that allows us to relate auditory model-based features with single trial perceptual outcomes. This allows us to test whether the foreground and background summary statistics outlined above can serve as explanatory acoustic features to predict speech recognition capabilities in the presence of arbitrary real-world environmental background sounds. Conceptually, we expect that the foreground and background sound power in the cochlear or mid-level auditory space can account for acoustic interference underlying speech recognition in noise perceptual abilities. To test this, we used the auditory model features from the cPS or mMPS as predictor variables for the GPR framework in order to predict digit recognition abilities in natural sources of environmental noise (Experiment 1 and 3).

The cPS and mMPS statistics described above, were first vectorized and concatenated for the background and foreground sounds yielding a composite feature vector, **X**, for each of the acoustic representations. Both the cPS and mMPS representations were tested separately, which allows us to test how each of the feature spaces contributes to perceptual interference. For instance, cPS representation contains only spectral cues and thus should be able to account for spectral interference between the foreground and background. By comparison, the mMPS representation also decomposes the sound into modulation components, and thus should be able to account for higher-level modulation interference. The composite feature vector for the cPS representation was obtained by concatenating cPS feature vectors from the foreground and background:

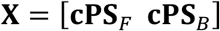

where the individual features in **CPS**_*F*_ and **CPS**_*B*_ are whitened so that they have unit variance and zero mean across experimental trials. Alternately, we also considered a mid-level feature representation. Here the composite feature vector was defined as

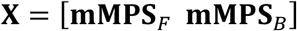

where **mMPS**_*F*_ and **mMPS**_*B*_ represent the vectorized foreground and background mid-level features. Given the high dimensionality of the cPS (57×2=114) and mMPS (57×7×17×2=13,566) spaces, principal component analysis is used to define dimensionality reduced feature vectors

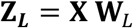

where **Z**_***L***_ are the principal component scores and **W**_*L*_ is an L-rank matrix containing the singular values of the covariance matrix, **X**^***T***^**X**. This dimensionality reduced representation is then used to fit a low-dimensional logistic regression model, which minimizes the possibility of overfitting. For experiment three, where we also varied the noise level, SNR was appended to **Z**_***L***_ as an independent explanatory variable to account for accuracy differences driven by the listening noise level. Here the probability of a correct behavioral decision is modeled by the logistic function

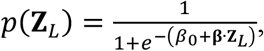

where *β*_0_ is regression intercept and the vector ***β*** = [*β*_1_ … *β*_*L*_] contains the regression weight parameters. For a given feature representation (cPS or mMPS), the regression weights were derived using cross validated maximum likelihood estimation, where we optimized for the number of explanatory variables by adjusting *L* between 1 to 100. For each iteration of *L*, the optimal regression weights were derived using the dimensionality reduced feature vector, **Z**_***L***_, as input to the model. The training and validation data samples were split using 25-fold cross validation and the cross-validated log-likelihood was estimated for each iteration of *L*. The optimal model dimensionality (*L*_*opt*_) was then chosen by selecting the number of principal components that maximized the cross-validated log-likelihood (**Figure S2**). Although the original stimulus dimensionality was substantially higher for the mid-level representation, the optimal cochlear and mid-level models had a similar dimensionality (cochlear *L*_*opt*_=20; mid-level *L*_*opt*_=24).

