## Supplemental Data and Figures for "Interference of mid-level sound statistics underlie human speech recognition sensitivity in natural noise"

Figure 1-1. Eight-Speaker Babble

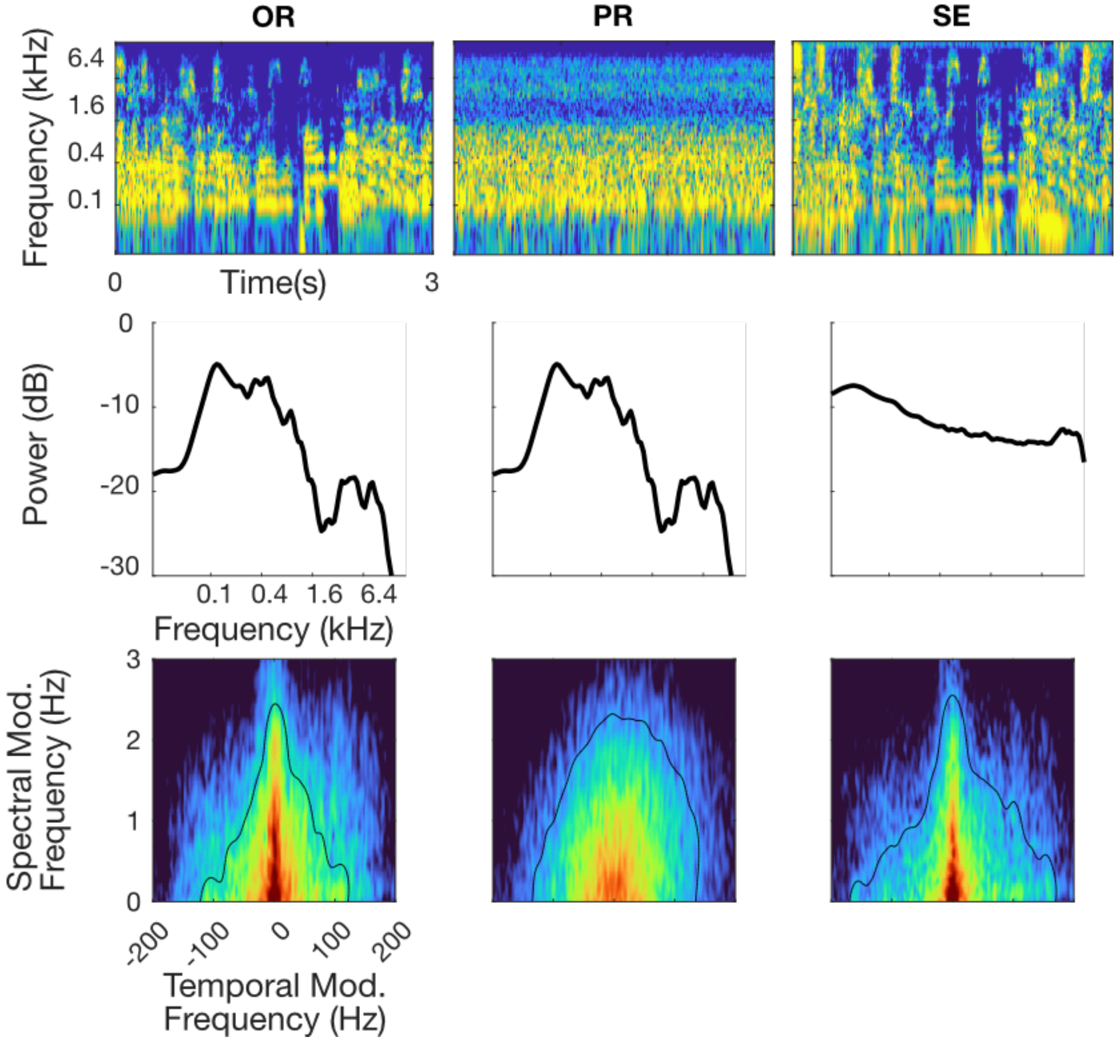


**Figure 1-1 to 1-10: Cochlear and modulation statistics for 3-second exemplars of all the backgrounds used in Experiment 1.** Each figure is formatted as for Fig. 1B. Each of the backgrounds is shown for the original (OR), phase randomized (PR), and spectrum equalize (SE) configurations (column 1-3, respectively). Cochleograms, cochlear power spectrum (cPS), and modulation power spectrum are shown in the 1$^{st}$, 2$^{nd}$ and 3$^{rd}$ rows. Note that regardless of the background, the PR condition preserves the original (OR) spectrum of the sound (middle panel, 2$^{nd}$ row) but whitens the modulation power (middle panel, 3$^{rd}$ row). Alternately, the SE condition whitens the sound cPS, but retains much of the original (OR) sound modulation content.

Figure 1-2. Factory

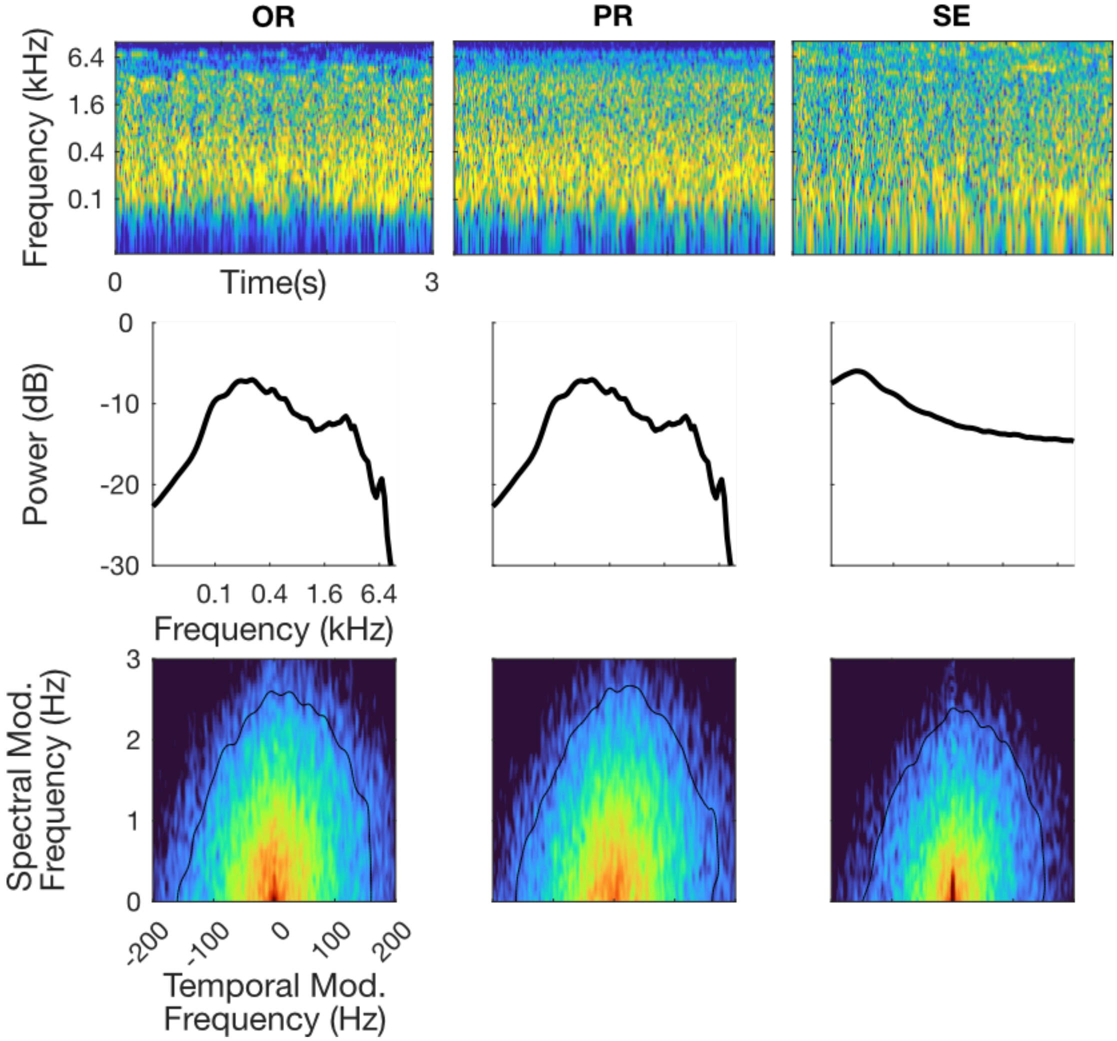

Figure 1-3. City

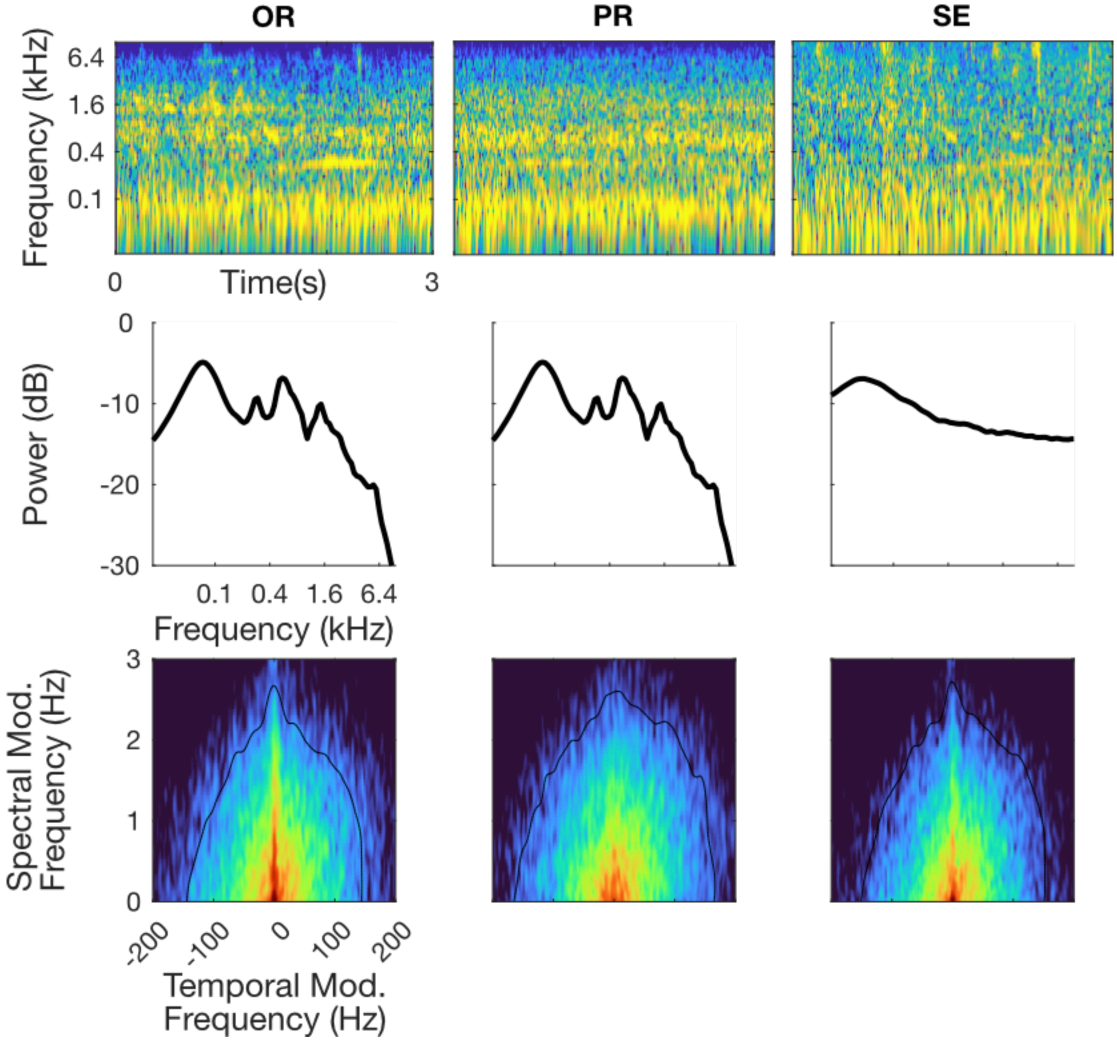

Figure 1-4. Two Speaker Babble

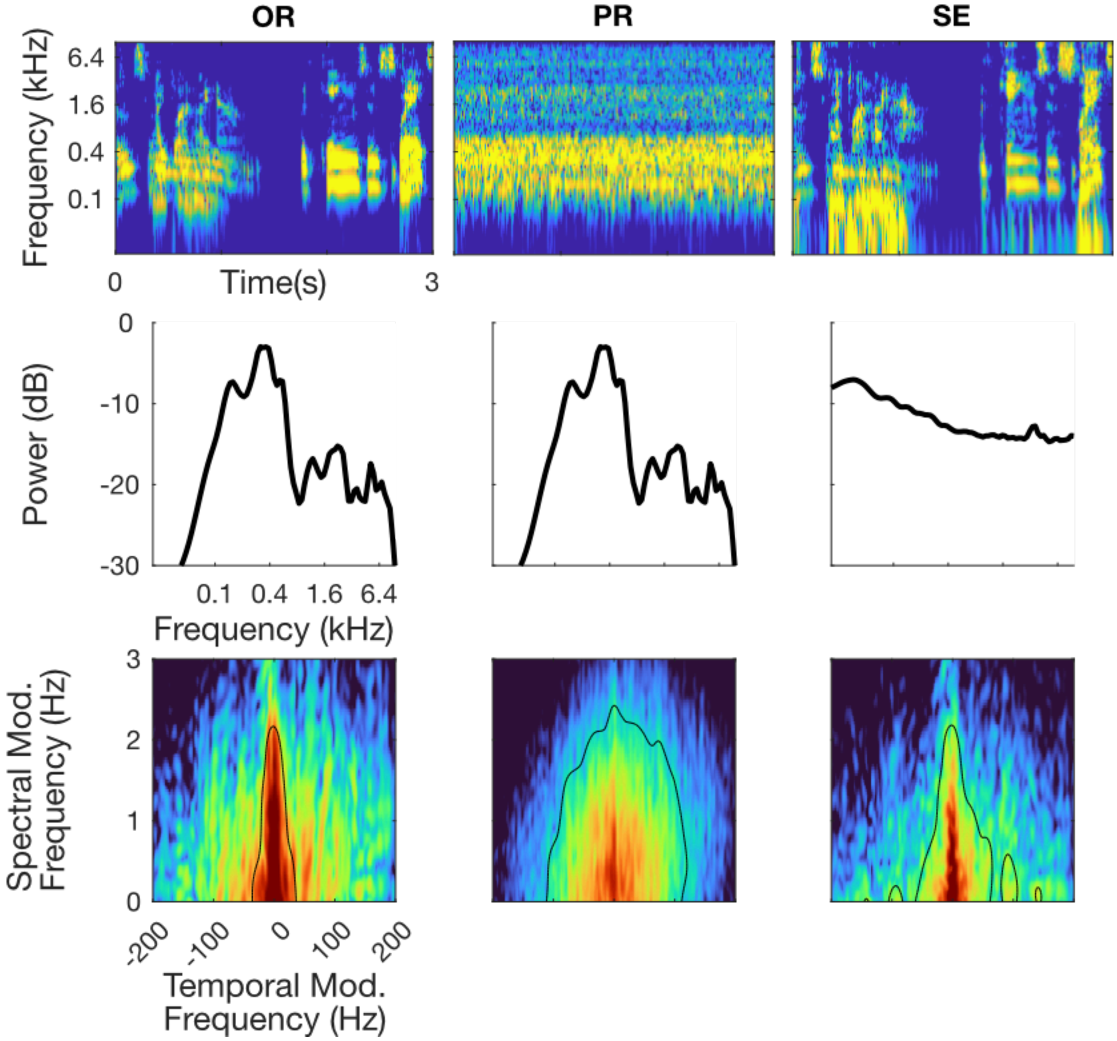

Figure 1-5. Water

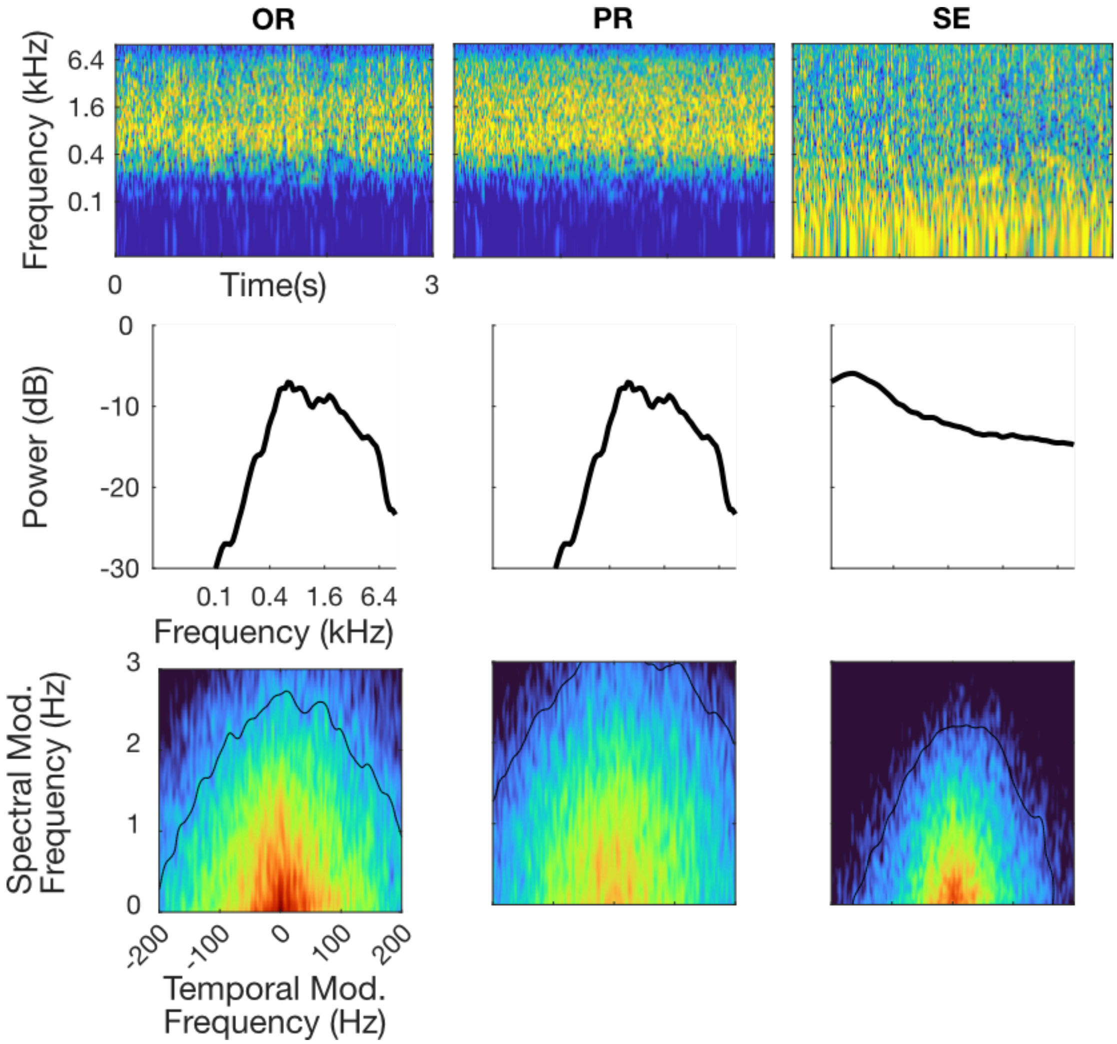

Figure 1-6. Construction

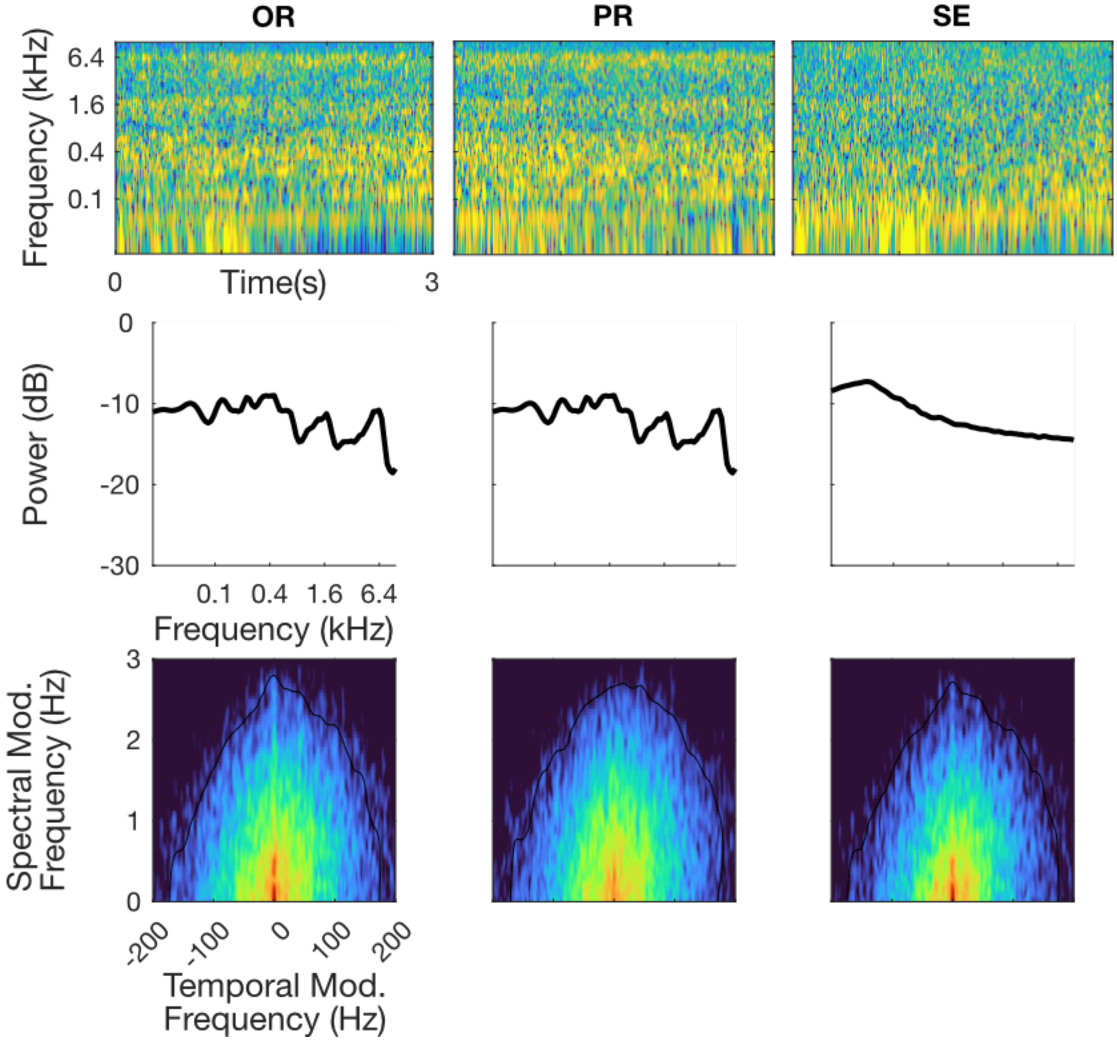

Figure 1-7. Jackhammer

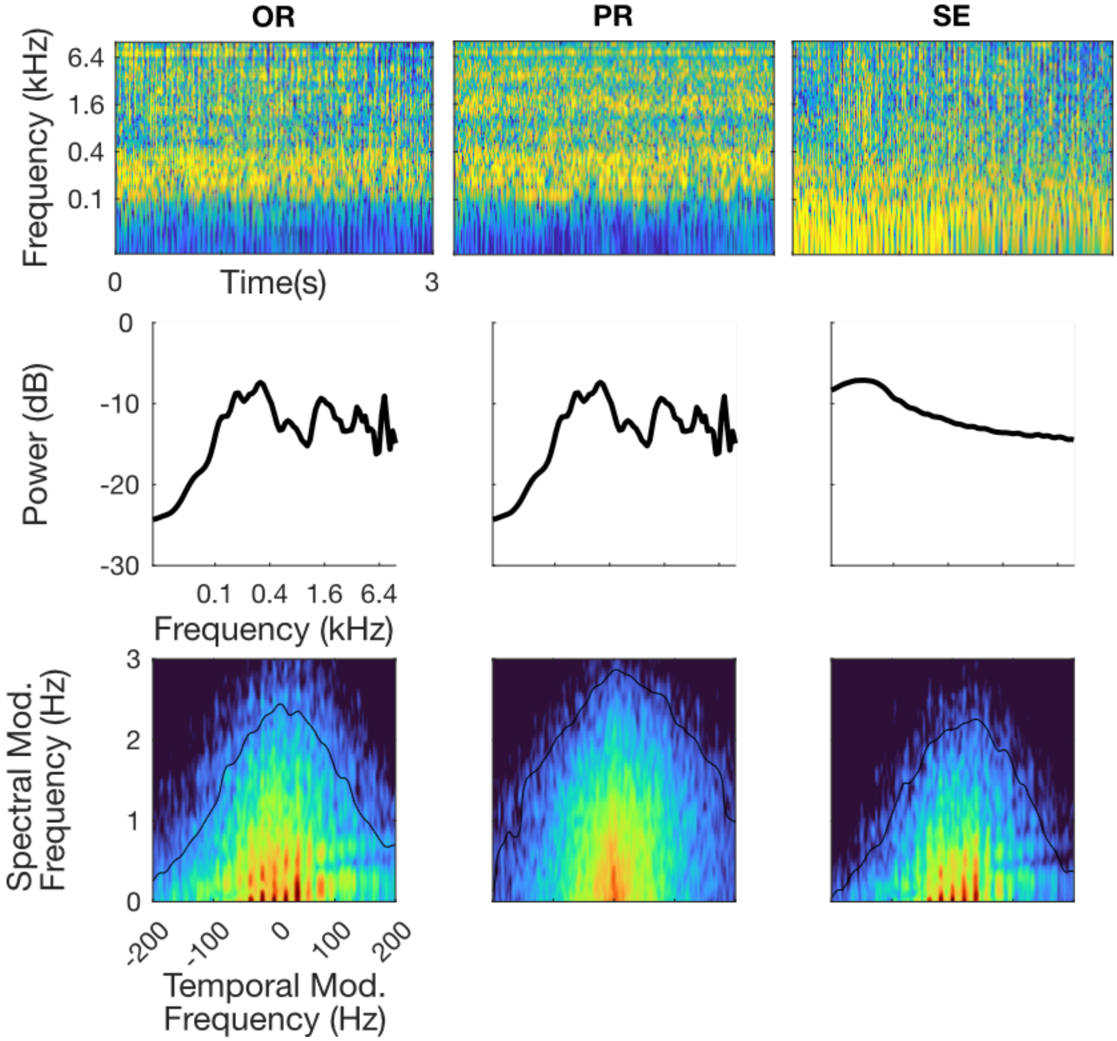

Figure 1-8. Freeway

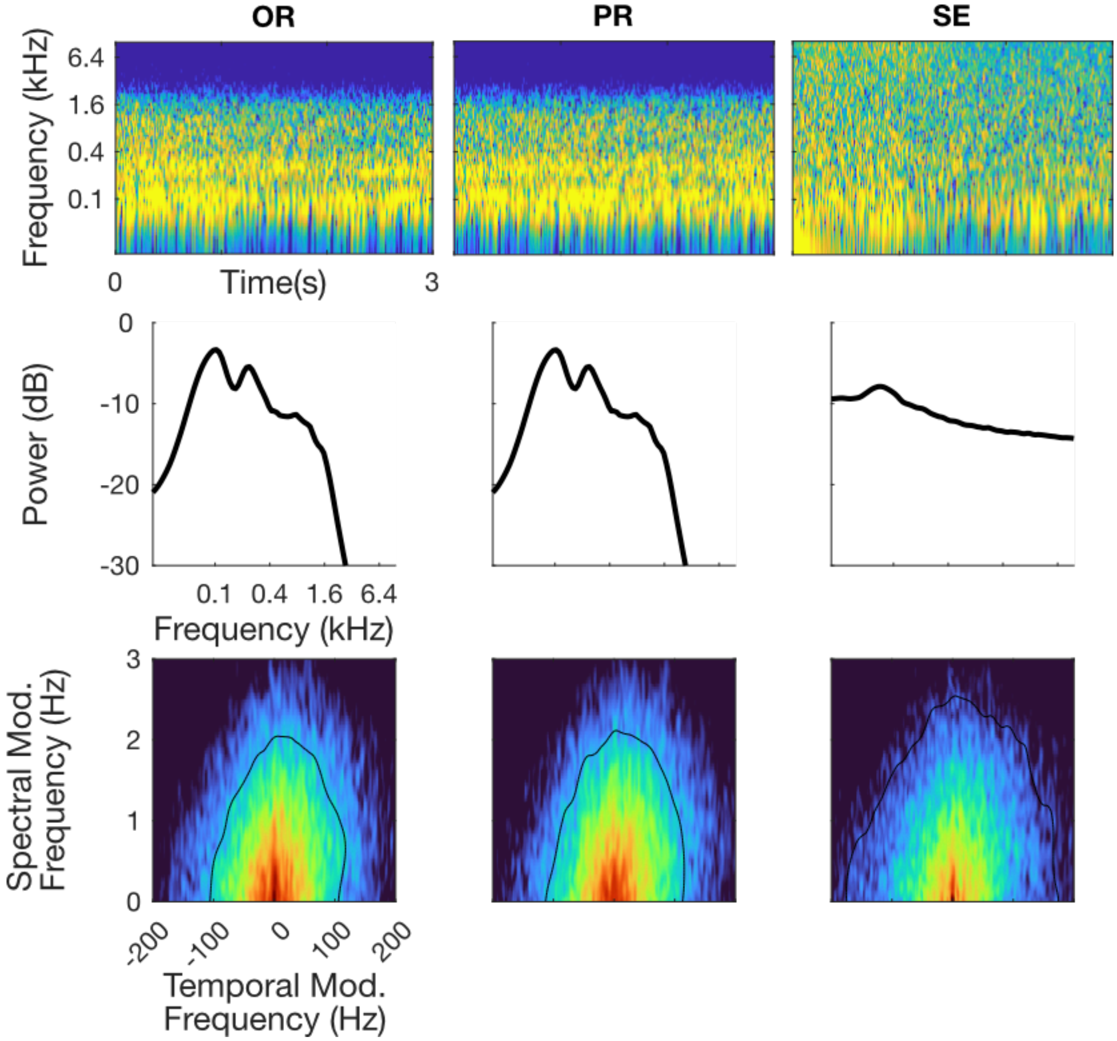

Figure 1-9. Bird Babble

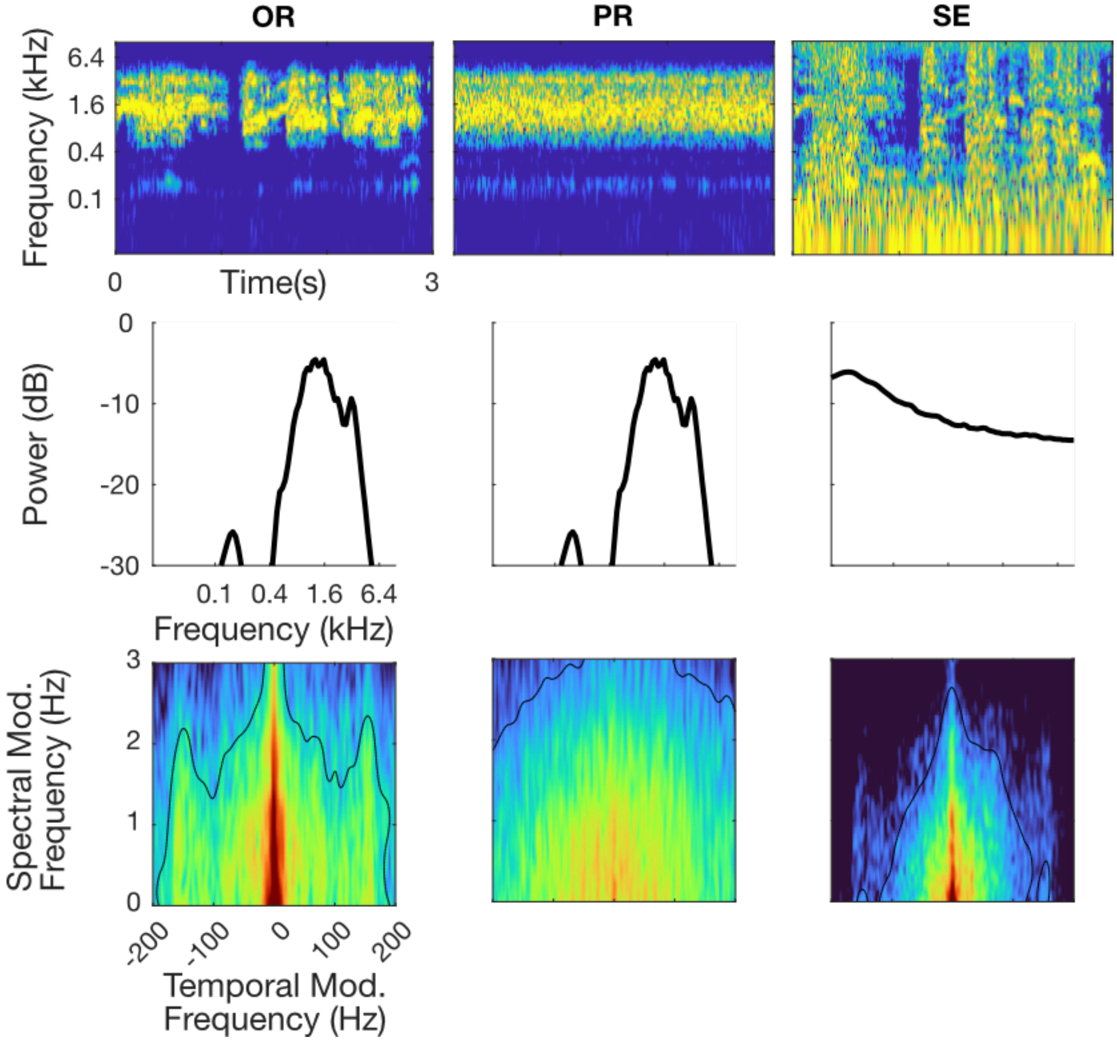

Figure 1-10. Fire

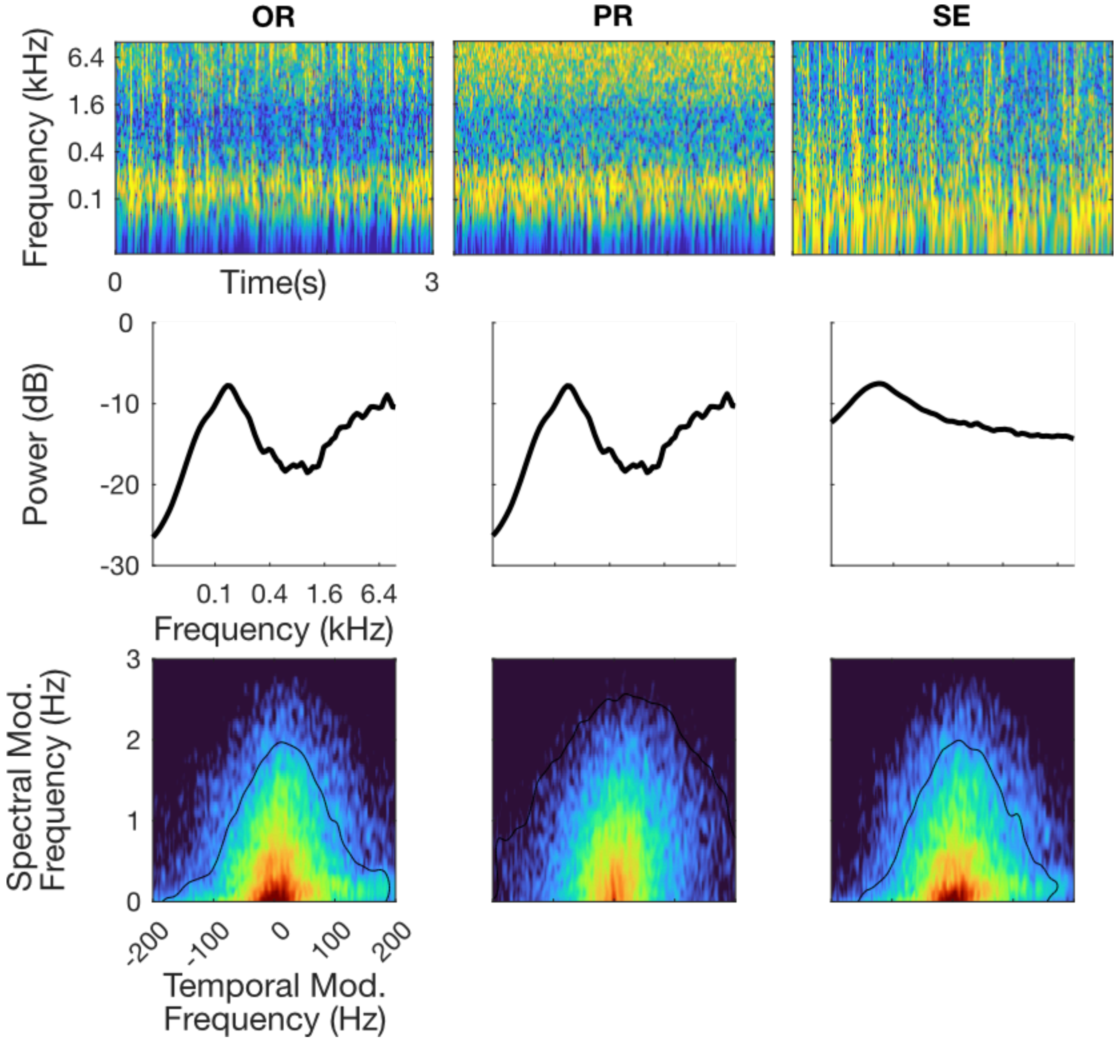

Figure 1-11. White Noise

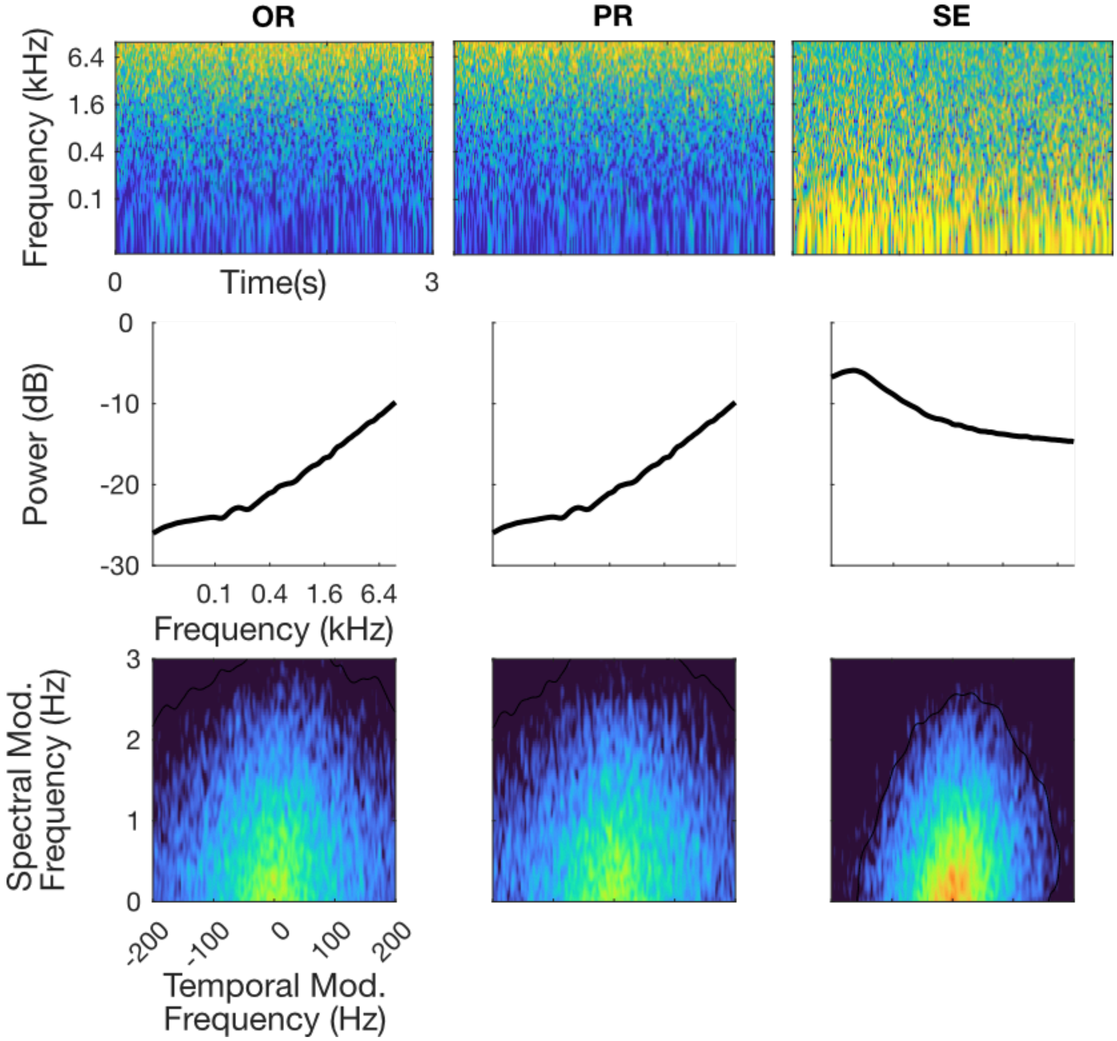

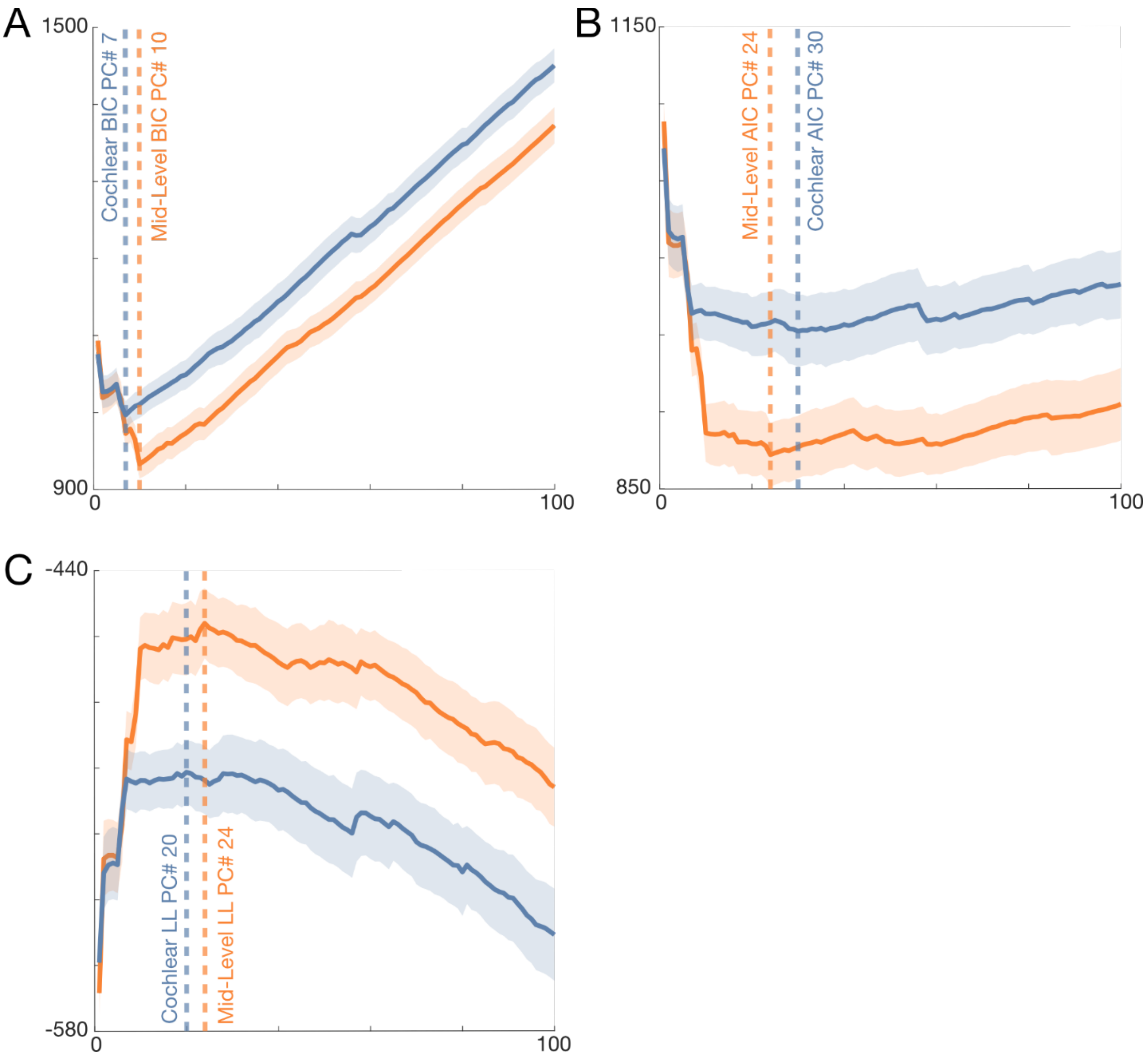


**Figure 4-1: Deriving the optimal number of principal components (PCs) for both the cochlear and mid-level GPR models.** For both the cochlear (blue) and mid-level GPR model (orange), we measured the (A) Bayesian information criterion (BIC), (B) Akaike information criterion (AIC), and (C) log-likelihood (LL) while varying the number of principal components used for each model. The curves indicate the average value of each metric and the shaded region indicates SD (across participants). Vertical dashed lines indicate the number of PCs needed to minimize the AIC or BIC or maximize the LL.

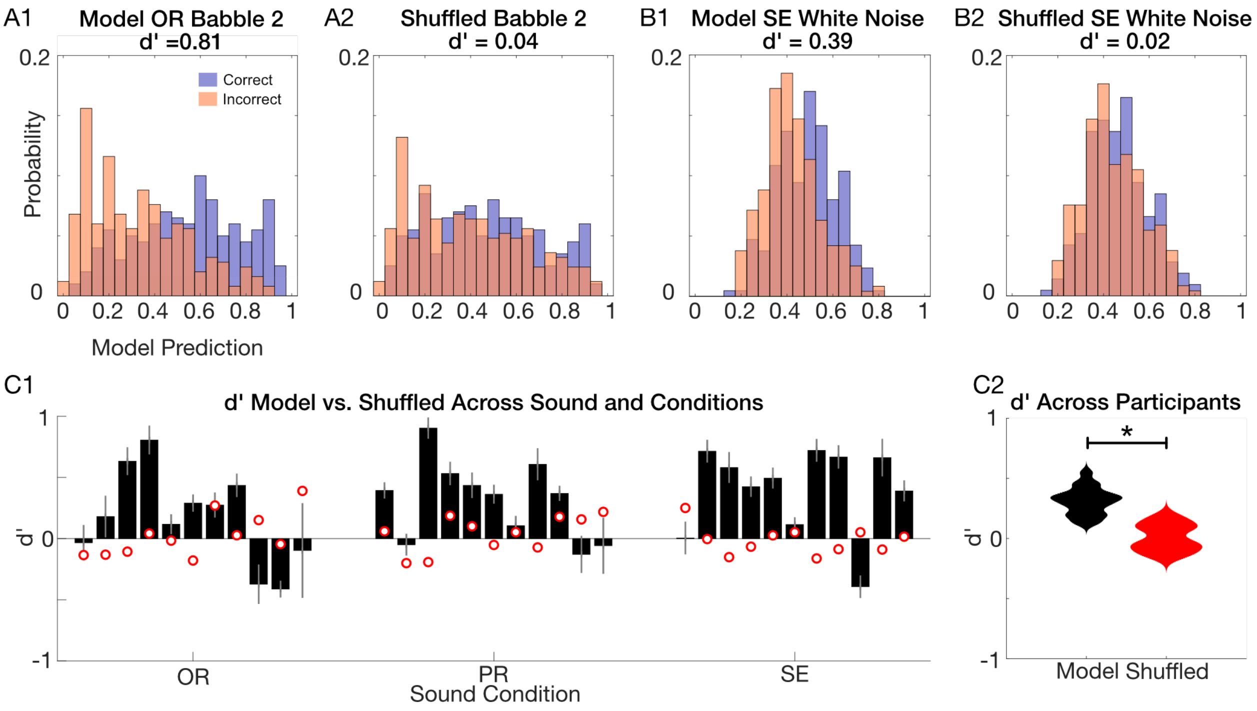


**Figure 4-2: Mid-Level GPR model accounts for single-trial changes in accuracy and outperforms a category-specific Bernoulli model for single trial predictions.** (A) Distribution of predicted accuracy for the mid-level model and category-specific Bernoulli model simulation (trial shuffled predictions) for two different backgrounds (A1-A2, Babble-2; B1-B2, SE White Noise). For both backgrounds, model predictions for correct trials (blue) were biased towards higher predicted accuracy while incorrect trials (pink) showed lower predicted accuracy, which for both cases produced a positive sensitivity index (A1, babble-2, d'=0.81; B1, SE White Noise, d'=0.39). When the prediction trials are shuffled (A2, B2), any association between the spectrotemporal sound cues, participant responses, and predicted performance is lost as expected for a Bernoulli process where correct and incorrect trials would occur independently of the acoustic cues for each background category. After trial shuffling, the model produces overlapping prediction distributions for correct and incorrect decisions and a low sensitivity index near zero (A2, babble-2, d'=0.04; B1, SE White Noise, d'=0.02). (C1) Histogram showing the sensitivity index for all the background sounds and different background conditions (OR, PR and SE). The background sounds are ordered in ascending OR accuracy (as in Fig. 1C). Bar plots show the average sensitivity index (*d*') for each background and condition for the Mid-Level GPR-model whereas the red dots show the sensitivity index (*d*') for the category-specific Bernoulli model (simulated using trial shuffling). Error bars designate the SEM (across participants). (C) Distribution of average sensitivity index (n=18 participants; averaged across all backgrounds and conditions, OR, SE, PR) for the mid-level model (black) and the category-specific Bernoulli model (trial shuffled simulation, red). * indicates a significant difference (t-test, $t(17)=7.9$, $p<10^{-6}$).

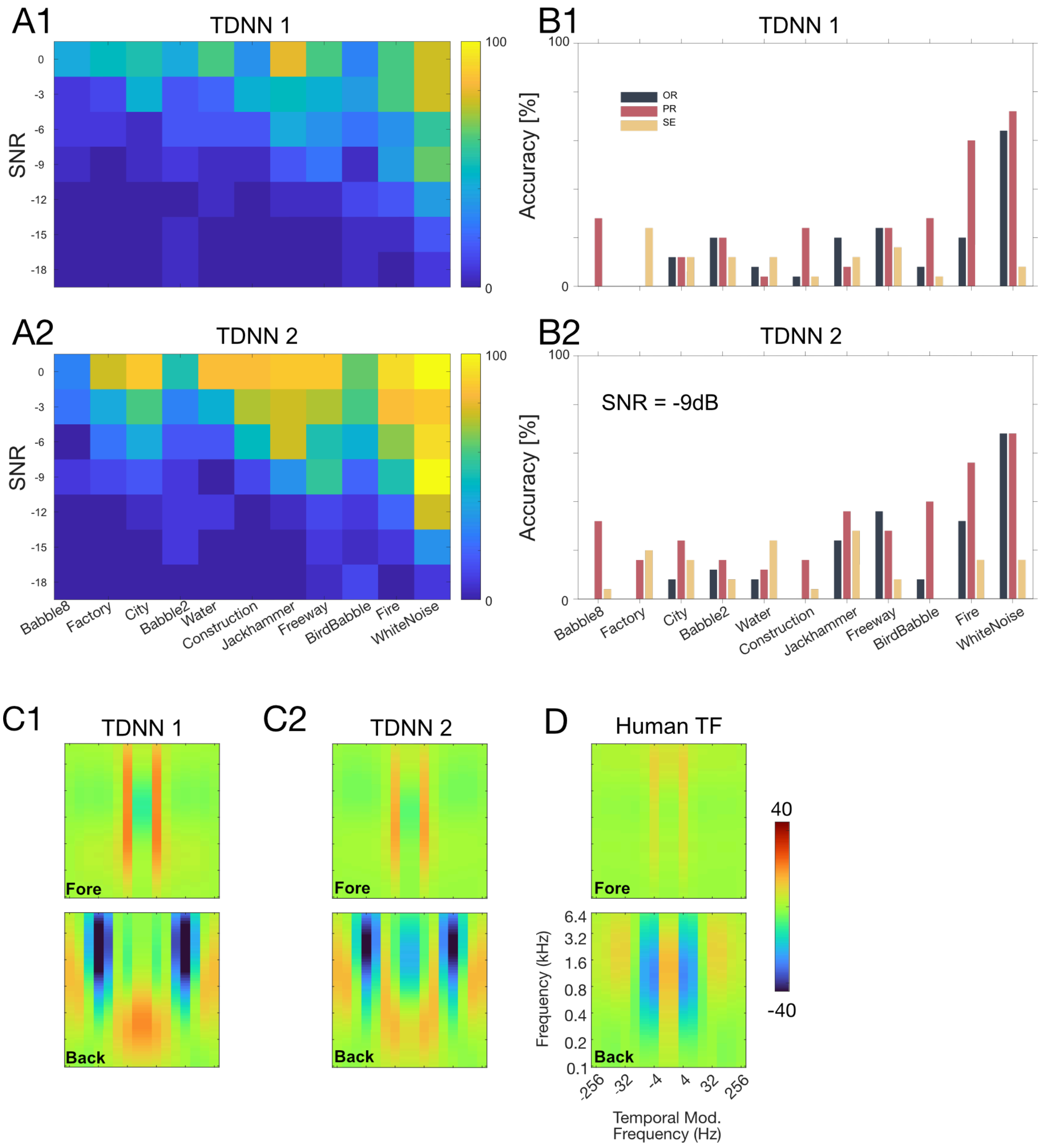


**Figure 4-3: Accuracy and GPR transfer functions for two neural network models diverges from human results.** Results are shown for the multiple SNR stimuli (as in Fig. 5; left, A1 and B1) and the OR/PR/SE stimuli at -9dB (as in Fig. 1; right A2 and B2). Here we tested task performance using two pretrained time delay neural network models (TDNN) with the pytorch-based SpeechBrain library (*6*). One network was trained for word recognition using speech commands (Google Speech Commands dataset (*7*)) and the second was trained for speaker recognition (Voxceleb1+ Voxceleb2 datasets) (*8-10*). Both networks generate low-dimensional embeddings (512-d) of variable length sound input using X-Vectors (*11*), and here we trained an additional readout layer (L2-regularized logistic regression) to identify the single digits 0-9 using our digit-in-noise stimuli. To make the model output comparable to the human data, results here show cross-validated three-digit accuracy calculated by combining model outputs across single digits. Note that unlike the human participants, the models have information about the exact start and stop times of the

foreground digits. As for the human data, the outputs of these models were fitted to our mid-level GPR model (using 30 PCs) and the model weights were used for model evaluation. (D) For reference, the average background and foreground GPR model human weights are shown for all of our participants. (C1-C2) The foreground and background weights of both networks are quite different from those of human listeners indicating that they employ different invariances and use different spectrotemporal cues that impact their performance. For instance, there is a strong background suppressive component ~32 Hz in both networks as well as positive hot-spots not visible for human listeners. Furthermore, the background weights of both networks are stronger than humans (shown on identical color scales) which is consistent with their lower digit recognition performance.

**Audio 1-1**

Example audio excerpts for each of the 33 background conditions for Experiment 1 are available at https://doi.org/10.12751/g-node.e7vt7m under the directory /SupplementalAudio/Audio_1/. All of the three-digit sequences are delivered at an SNR -9 dB. Exemplars for each of the 11 background sound conditions are organized in a directory corresponding to the experimental manipulation performed (OR, PR, SE). Sounds are numbered from 1 to 11 in rank order according to the perceptual accuracy, as in Fig. 1C (Sound01.wav, Sound02.wav, etc.).

**Audio 2-1**

Example audio excerpts for each of the background conditions for Experiment 2 are available at https://doi.org/10.12751/g-node.e7vt7m under the directory /SupplementalAudio/Audio_2/. The background conditions include the babble-8 and jackhammer backgrounds delivered at variable SNR and variable statistics (Spec, +Mar, +MPS, +Corr, and Orig). Sounds are organized in subdirectories corresponding to the SNR condition (-12, -9, -6, and -3 dB) and background name (EightSpeakerBabble or Jackhammer) and are labeled according to the summary statistics used to synthesize the background (SoundSpec.wav, SoundMar.wav, SoundMPS.wav, SoundCorr.wav, SoundOrig.wav).
